# Interactive effects of microplastic pollution and heat stress on reef-building corals

**DOI:** 10.1101/2021.07.30.454458

**Authors:** Jessica Reichert, Vanessa Tirpitz, Rajshree Anand, Katharina Bach, Jonas Knopp, Patrick Schubert, Thomas Wilke, Maren Ziegler

## Abstract

Plastic pollution is an emerging stressor that increases pressure on ecosystems such as coral reefs that are already challenged by climate change. However, the effect of plastic pollution in combination with global warming is largely unknown. Thus, the goal of this study was to determine the cumulative effect of microplastic pollution with that of global warming on reef-building coral species and to compare the severity of both stressors. For this, we conducted a series of three controlled laboratory experiments and exposed a broad range of coral species (*Acropora muricata, Montipora digitata, Porites lutea, Pocillopora verrucosa*, and *Stylophora pistillata*) to microplastic particles in a range of concentrations (2.5–2,500 particles L^-1^) and mixtures (from different industrial sectors) at ambient temperatures and in combination with heat stress. We show that microplastic can occasionally have a negative effect on the corals’ thermal tolerance. In comparison to heat stress, however, microplastic constitutes a minor stressor. While heat stress led to decreased photosynthetic efficiency of algal symbionts, and increased bleaching, tissue necrosis, and mortality, treatment with microplastic particles had only minor effects on the physiology and health of the tested coral species at ambient temperatures. These findings underline that while efforts to reduce plastic pollution should continue, they should not replace more urgent efforts to halt global warming, which are immediately needed to preserve remaining coral reef ecosystems.

**Graphical Abstract:** 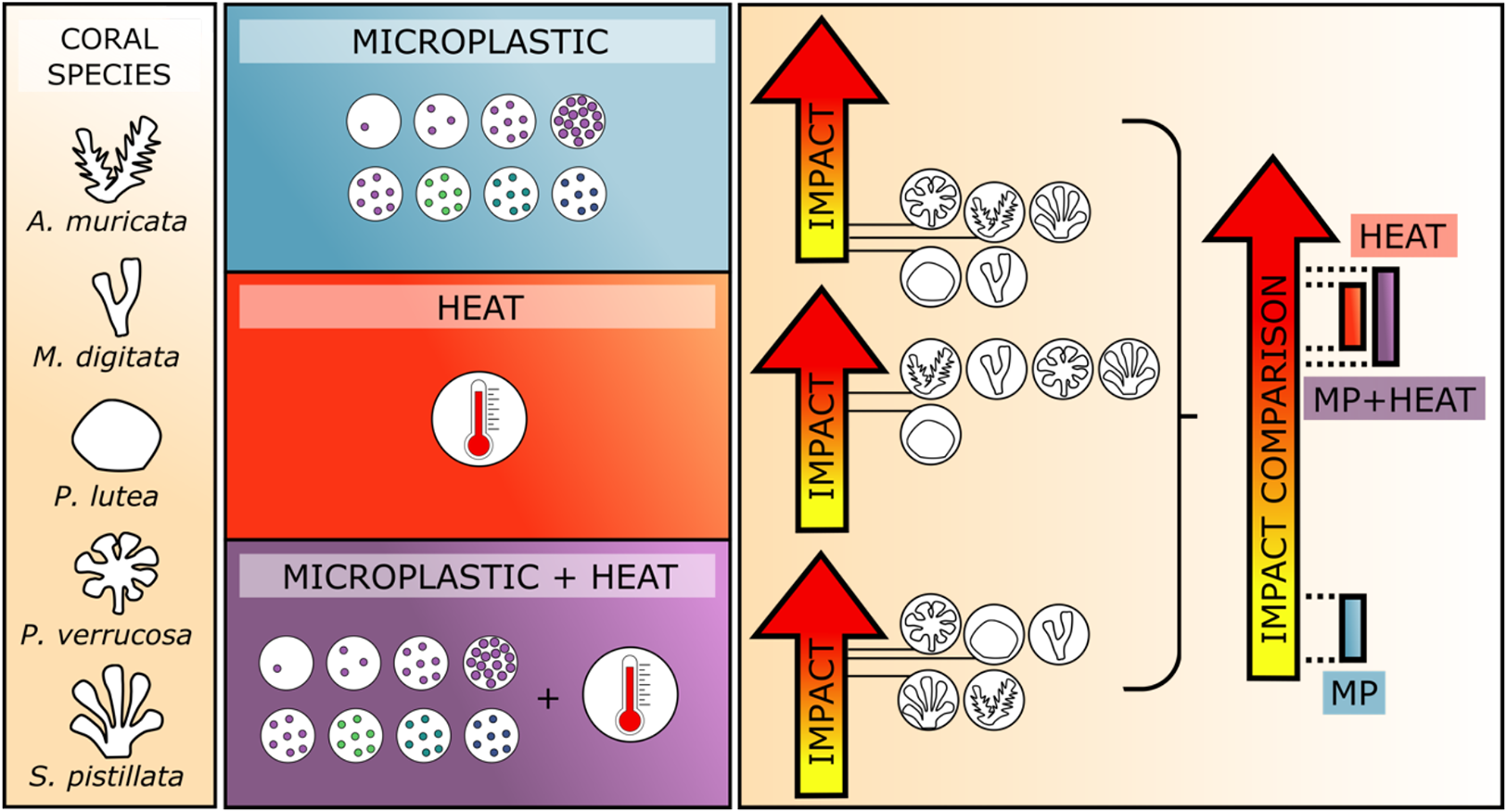

## Introduction

Coral reefs provide important ecosystem services such as shoreline protection, subsistence for hundreds of million people, and a habitat for a diverse range of organisms (Fisher et al., 2015). Yet, worldwide coral reefs are declining and mass mortalities of reef-building corals are mainly driven by recurrent marine heat waves as a consequence of global climate change (Frölicher et al., 2018; Hughes et al., 2018). Heat stress disrupts the obligate symbiosis of reef-building corals with dinoflagellate microalgae (Symbiodiniaceae) and causes coral bleaching and widespread coral mortality (Hoegh-Guldberg, 1999; Hughes et al., 2018). Also, a range of local stressors such as nutrient enrichment and pollution of coastal waters are often linked with a significant loss of coral cover and diversity, and have been shown to decrease the tolerance of reef-building corals to heat stress (Wiedenmann et al., 2013).

The accelerating pollution of coastal waters with plastic is an emerging stressor that affects reef-building corals worldwide (Lamb et al., 2018). Especially microplastic (MP; i.e., particles < 1 mm; Hartmann et al., 2019) is suspected to pose an additional threat to corals. Microplastic occurs in various shapes (most often in form of particles and fibers), polymer types (most commonly polyethylene and polypropylene) and highly variable concentrations spanning eight orders of magnitude from less than 0.0001 particles per L (Jensen et al., 2019) to more than 10,000 particles per L (Badylak et al., 2021). The effect of microplastic on corals ranges from low and moderate (Chapron et al., 2018; Mouchi et al., 2019; Reichert et al., 2019) to severe (Syakti et al., 2019; Tang et al., 2018) depending on the experimental approach and particle dose used. Corals actively ingest (Allen et al., 2017; Hall et al., 2015; Reichert et al., 2019) or passively catch particles by adherence (Corona et al., 2020; Martin et al., 2019). This has been shown to interfere with the feeding performance of the corals (Mouchi et al., 2019; Savinelli et al., 2020). Additionally, leachates (e.g., phthalates) are likely transferred from the particles to the corals (Montano et al., 2020; Saliu et al., 2019). Corals respond with changes in photosynthetic performance of the associated symbionts (Lanctôt et al., 2020; Reichert et al., 2019; Su et al., 2020), alterations in metabolite profiles (Lanctôt et al., 2020) and stress enzyme activity (Tang et al., 2021, 2018), reduced growth (Chapron et al., 2018; Reichert et al., 2019), and decreased overall health (Reichert et al., 2019; Syakti et al., 2019).

Seeing these impacts, the motivation to reduce plastic pollution in the oceans is high and the public perceives plastic pollution as the greatest threat, far worse than global warming (Stafford and Jones, 2019). Although this attention creates direct and much-needed incentives to counteract the accelerating plastic problem, it remains unclear to which extent resources should be divided between climate change mitigation and reduction of plastic pollution. This is mainly because the impact of plastic pollution on coral reefs in a warming ocean is poorly understood. Therefore, integrative research approaches to assess the combined effects of plastic pollution and global warming on coral reef ecosystems in the Anthropocene are needed.

Indeed, plastic pollution holds a high potential to amplify the effects of global warming and might decrease the thermal tolerance of reef-building corals, as seen for other additive stressors (Wiedenmann et al., 2013). Contact with and uptake of plastic particles may inhibit feeding and possibly lead to a depletion of energy reserves in corals (Chapron et al., 2018). This may be critical as the survival and recovery of corals under heat stress is, in part, dependent on their energy reserves (Grottoli et al., 2006). In addition, the virulence of some pathogenic bacteria increases with temperature (Konkel and Tilly, 2000). Microplastic particles may act as vectors for such pathogenic bacteria (Franco et al., 2020; Kirstein et al., 2016), which are taken up by coral polyps (Rotjan et al., 2019). However, while plastic leachates impair aquatic photosynthesis in phytoplankton (Tetu et al., 2019), the exposure of corals to MP leads to increased photosynthetic efficiency of the algal endosymbionts (Lanctôt et al., 2020; Reichert et al., 2019). In addition, it is unclear how these effects are modulated under heat stress.

Despite these potential interactions, the combined effect of MP and temperature stress on reef-building corals has not been investigated yet. Therefore, the goal of this study was to assess the effects of MP pollution on the thermal tolerance of reef-building corals in order to better relate the two stressors and guide future policy making. Specifically, we tested the effect of microplastic particles at ambient temperature and in combination with heat stress I) using different particle concentrations and targeting two reef-building coral species, II) and on a broader spectrum of major reef-building coral genera, targeting five coral species. Finally, we tested III) whether the observed effects are consistent across different mixtures of microplastic particles beyond a single polymer. We assessed the health and performance of the coral holobiont under separate and combined microplastic and heat stress. To this end, we measured the photosynthetic efficiency of the microalgal symbionts, coral bleaching intensity, host tissue necrosis, and mortality in three independent experiments (Fig. 1).

**Figure 1.**
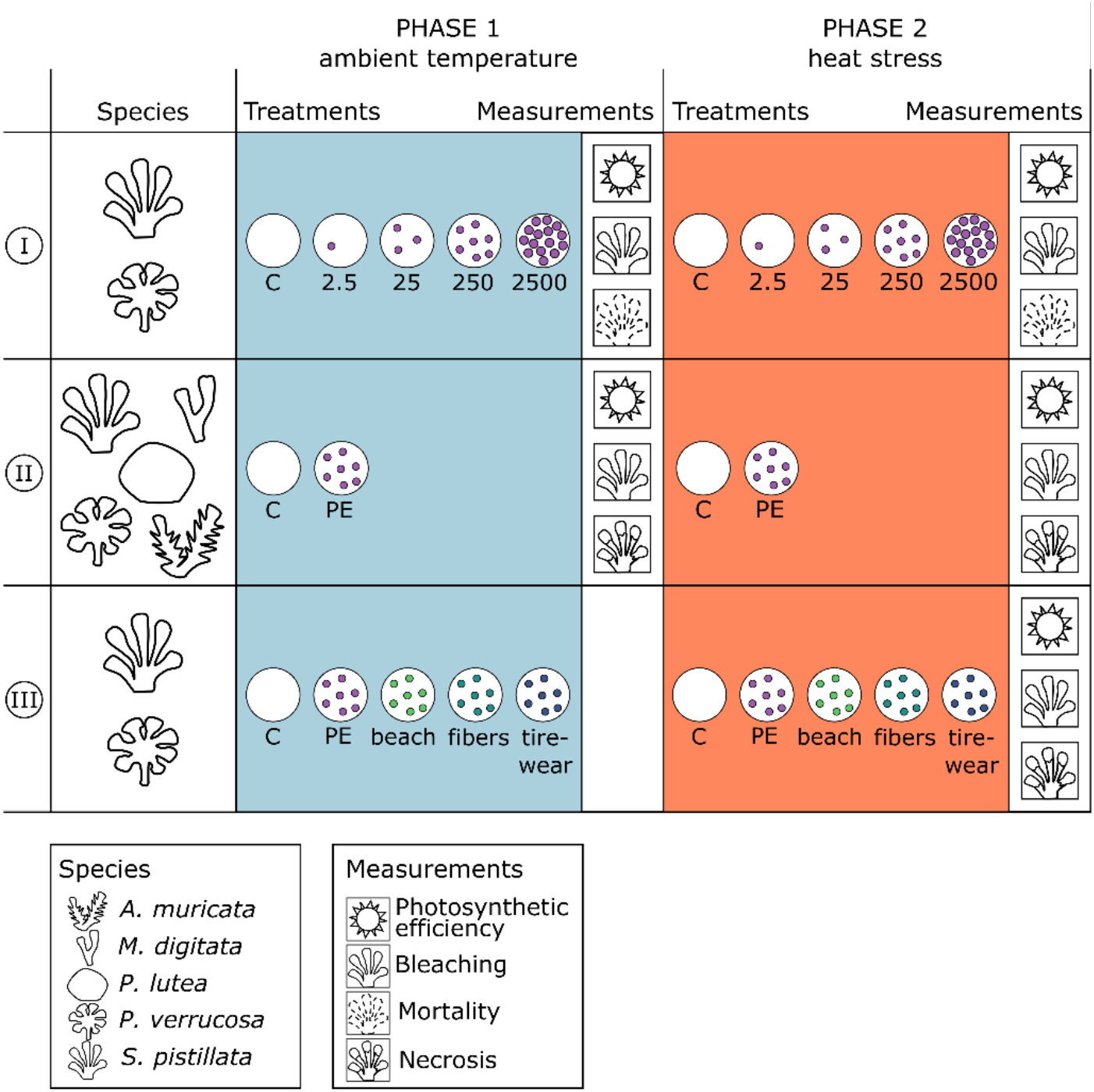
Study design and key parameters. Summary of three independent experiments testing the effects of microplastic and heat stress on coral health and photosynthetic performance.

## Materials and Methods

We conducted a series of three independent experiments to investigate the effect of microplastic pollution on heat stress tolerance of stony corals between September 2018 and November 2019. All experiments took place in the ‘Ocean2100’ aquarium facility of the Justus Liebig University Giessen, Germany.

### Experiment 1 – Effect of different concentrations of microplastic on coral heat stress tolerance

The first experiment was designed to investigate the effect of different concentrations of microplastic particles on coral heat stress tolerance and thus to determine whether current or projected future microplastic concentrations have an effect on the performance of corals under climate change conditions. The experiment was comprised of five treatments with increasing target concentrations of polyethylene (PE) microplastic particles over nine weeks (Tab. S1). For this, we added 0, 4, 40, 400, or 4,000 mg to each of three replicate tanks (0.1–100 mg L^-1^, respectively) resulting in 0 (microplastic-free), 2.5, 25, 250, and 2,500 particles L^-1^ in the water column, respectively (Fig. S1). This range of concentrations spans values that have already been observed in coral reef environments (0.00012–126 particles L^-1^ (Jensen et al., 2019; Patterson et al., 2020)). It extends over future projected concentrations (250 particles L^-1^) that may be expected for polluted oceans, based on the estimated three-fold increase in plastic pollution until 2100 (Koelmans et al., 2017), and up to very high values (2,500 particles L^-1^) that we used to exert an extreme stress on the coral fragments. After six weeks of microplastic treatment, water temperatures were increased in all tanks (microplastic and microplastic-free controls) over one week by 1 °C day^-1^ from 27 to 33 °C and held at 33 °C for another two weeks, mimicking heat stress conditions in coral reefs (Ainsworth et al., 2016; Fig. S2, Tab. S1).

For this experiment, we focused on two common reef-building coral species, *Pocillopora verrucosa* (Ellis & Solander, 1786) and *Stylophora pistillata* (Esper, 1797). *P. verrucosa* is known to frequently react to microplastic (Reichert et al., 2018) and *S. pistillata* is a commonly investigated coral model species. Eight source colonies per species were each cut into 15 fragments, which were suspended on nylon strings, approximately 5 cm below the water surface (Fig. S3). Fragments from each colony were present in each of the 15 experimental tanks (three replicate tanks per treatment, n = 24 fragments per treatment).

To monitor the physiological condition of the coral fragments in response to exposure to increasing concentrations of microplastic and additional heat stress, we measured the light-adapted (ΔF/Fm’) and dark-adapted (Fv/Fm) photosynthetic efficiency at the beginning of the experiment, after five weeks during the MP phase, and a last time during the heat-stress phase (Tab. S2), which was determined for each colony separately by at least three fragments with > 50 % bleached or necrotic tissue. At this last time point, we also assessed the thermal bleaching response with standardized photographs (Tab. S3, see below for details). Subsequently, fragments were returned to the tanks for survival analyses and scored twice a day (10:00 and 18:00 h) for remaining tissue. Fragments were recorded as dead, when they had lost all tissue from the skeleton.

### Experiment 2 – Effect of microplastic on heat tolerance of different stony coral species

The second experiment was designed to investigate the combined effect of microplastic pollution and heat stress on a broader selection of common reef-building coral species. We exposed five stony coral species to PE microplastic particles at 230 particles L^-1^ (i.e., 400 mg tank^-1^ / 10 mg L^-1^, Fig. S4) over 11 weeks (Tab. S1). For this, we mounted six fragments from each of three coral colonies per species (*Acropora muricata* (Linnaeus, 1758), *Montipora digitata* (Dana, 1846), *Porites cylindrica* (Dana, 1846), *P. verrucosa, S. pistillata*) onto self-made concrete bases (Fig. S5, Tab. S4). Coral fragments were distributed over six experimental tanks, so that one fragment from each colony was present in each tank (three microplastic tanks (n = 9) and three microplastic-free tanks (n = 9), respectively). After five weeks of microplastic treatment, water temperatures were increased in all tanks (microplastic and microplastic-free controls) over two weeks by 0.5 °C day^-1^ from 26 to 30 °C and then by 0.25 °C day^-1^ to the final holding temperature of 32 °C, which was held for another three weeks (Fig. S6, Tab. S1). This design followed naturally occurring temperature trajectories in coral reefs (Ainsworth et al., 2016).

To monitor the physiological condition of the coral fragments in response to microplastic exposure and additional heat stress, we measured the light-adapted (ΔF/Fm’) and dark-adapted (Fv/Fm) photosynthetic efficiency nine times at roughly weekly intervals from the start of the experiment until values dropped below 0.3 (Tab. S2). To assess the thermal bleaching response of the coral fragments, standardized photographs of each fragment were taken at four time points weekly, starting with the increase of experimental temperatures (Tab. S3). These pictures were used to calculate tissue brightness as a measure of coral bleaching. Because tissue necrosis (i.e., loss of tissue, indicated by a sharp edge of remaining tissue in contrast to the blank skeleton) is a common response of stony corals to microplastic exposure, we scanned each fragment with a hand-held 3D scanner to create three-dimensional models of the surface area and determine the fraction of necrotic surface. Surface scans were obtained at the end of the experiment, which was determined for each colony separately, if at least one fragment of that colony had > 50 % bleached or necrotic tissue.

### Experiment 3 – Effect of different mixtures of microplastic particles on coral heat stress tolerance

The third experiment was designed to investigate the effect of different mixtures of microplastic particles on coral heat stress tolerance. This is important, as microdebris in the oceans is composed of a complex mix of manufactured and modified materials, with plastics, especially in the form of fibers, generally constituting the most common items in coral reef environments (Carr, 2017; Kroon et al., 2018). The experiment was comprised of five treatments over ten weeks (Tab. S1), which are representative of different major microplastic-producing industrial sectors: fashion industry with artificial clothing fibers (“fibers”), automobile sector with tire wear, brake abrasion, and varnish (“tirewear”), secondary marine microplastic from the beach (“beach”), the reference PE particles used in the first and second experiment (“PE”), and a microplastic-free treatment (“control”) (see below for detailed characterization of microplastic). 400 mg (10 mg L^-1^) of each mix of microparticles was added to the respective tanks (n = 3), corresponding to 86 (± 63 sd) beach particles L^-1^, 265 (± 228 sd) fiber particles L^-1^, 66 (± 50 sd) tire wear particles L^-1^, 242 (± 126 sd) PE particles L^-1^ (Fig. S7, S8). After nine weeks of the microplastic treatment, water temperatures were repeatedly increased daily following a short-term heat stress design(Voolstra et al., 2020). Each day at 12:30 h water temperatures were increased in all tanks (microplastic and microplastic-free controls) from 27 °C (ambient temperature) to the daily target temperature in 3 h, held for 3 h, and returned to ambient in 1 h until 19:30 h. The daily target temperature of 33.5 °C was determined in trials before the experiment and increased by 0.5 °C each day up to 35 °C, after which the holding time was extended to 6 h for two further days of short-term heat stress, according to the response of the coral fragments in the experiment (Fig. S9).

For this experiment, we again used the two common reef-building coral species, *Pocillopora verrucosa* and *Stylophora pistillata*. Nine source colonies per species, were each cut into five fragments, which were suspended on nylon strings, approximately 5 cm below the water surface (Tab. S4). The 15 experimental tanks (three replicates per treatment) were split into three blocks of five tanks. Five fragments of every colony were distributed over the five treatments within a block, with the fragments from the respectively same three colonies in the same tanks (n = 9 per treatment).

To monitor the physiological condition of the coral fragments in response to exposure to microplastic and the additional short-term heat stress, we measured the light-adapted (ΔF/Fm’) and dark-adapted (Fv/Fm) photosynthetic efficiency at the end of the heat-stress phase (Tab. S2), which was determined for each colony separately by at least one fragment with > 50 % bleached or necrotic tissue. At this last time point, we also assessed the thermal bleaching response with standardized photographs (Tab. S3) and tissue necrosis with surface scans (see below for details).

### Characterization of microplastic particles

The coral fragments in the three heat stress experiments were exposed to different mixtures of microplastic particles. Particle sizes were chosen to be smaller than the corallites of the tested coral species (300-1400 µm, depending on the species, Veron, 2000). In each of the experiments, we used black high density polyethylene (PE) microplastic (density: 0.95 g cm^3^, Novoplastic, Germany), which is one of the most common types of plastic in the oceans (Andrady, 2017) and has previously been used in similar experimental designs (Reichert et al., 2019, 2018). PE particles had an irregular shape with a rough surface structure, resembling natural secondary microplastic (Fig. S8). We strained particles to a size between ∼100–355 µm, mean particle length was 370 µm ± 112 SD and mean width was 225 µm ± 69 SD. For investigating the effect of different mixtures of microplastic particles on coral heat stress tolerance (Experiment 3) we used the reference PE particles and three additional mixtures of particles, which are representative of different major microplastic-producing industrial sectors (Fig. S8). (A) Fashion industry (“fibers”): artificial clothing fibers aimed to represent a naturally occurring mixture of fibers (Browne et al., 2011), which is often the most abundant type of microplastic in coral reef environments (Carr, 2017; Kroon et al., 2018). We manually cut a mix of fibers from synthetic fabrics (mean length 1,006 µm ± 835 SD, mean width 16 µm ± 4 SD) and analyzed their composition using ATR-FTIR against reference spectra from Baseman Polymer reference Database (Primpke et al., 2018). The used fabrics were composed of approx. 70 % acrylic and 30 % polyamide, or of 100 % polyester (Fig. S10). We mixed the resulting fibers to represent a nature-oriented compilation of 67 % polyester, and approx. 23 % acrylic and 10 % polyamide (Browne et al., 2011). (B) Automobile sector (“tirewear”): a combination of residues from the automobile sector composed of tire wear, brake abrasion, and varnish. This mixture was retrieved from the residues found on a cart track (EKS-Kartcenter-Cologne GmbH, Germany). Carts at the track were equipped with hydraulic brake systems and wheels from Dunlop (Cart Tires, Goodyear Dunlop Tires Germany GmbH, Germany). Particles were sieved to a size between ∼100–355 µm, mean particle length was 583 µm ± 234 SD, mean particle width was 218 µm ± 71 SD. (C) Secondary marine microplastic from the beach (“beach”): For this, plastic debris was randomly collected in April and August 2016 from beaches on Texel, The Netherlands, and fragmented by cryogenic milling, resulting in a mix of 60.9 % polyethylene, 27.7 % polypropylene, 3.1 % polyamide, 1.9 % polyurethane, 1.7 % polystyrene, 0.9 % polyvinyl chloride, 0.5 % polyethylene terephthalate, 0.2 % Ethylene-vinyl acetate, 0.1 % Ethylene-vinyl acetate, and 3.1 % unidentified particles (% by mass), (Kühn et al., 2018). The particles were sieved to a particle size of > 100 µm, mean particle length was 346 µm ± 240 SD, mean particle width was 184 ± 126 SD. All particles were conditioned in seawater from the aquarium system for at least two days before the start of the experiment.

### Microplastic particle quantification

The concentrations of microplastic particles were determined for each tank at weekly intervals and new particles were added when concentration decreased over time (due to tank maintenance activities, or ingestion and overgrowth by coral fragments). For particle quantification, we vacuum filtered three replicates of 100 ml of water per tank onto a cellulose filter with a mesh size of 8 µm (Whatman filter papers Grade 540, General Electric Healthcare Life Sciences, USA). In Experiment 1, 10 ml were filtered in the highest concentration (2,500 particles L^-1^). Particles on the filters were counted under a stereomicroscope and numbers extrapolated to calculate the concentration per liter.

### Coral species and fragmentation

We investigated the heat stress tolerance of a range of stony coral species under microplastic exposure. The studied species include: *Acropora muricata* (Linnaeus, 1758), *Montipora digitata* (Dana, 1846), *Porites cylindrica* (Dana, 1846), *Pocillopora verrucosa* (Ellis & Solander, 1786), and *Stylophora pistillata* (Esper, 1797). For Experiment 2, all coral colonies originated from the Indopacific and were sourced from the coral aquarium facility ‘Ocean2100’ of the Justus Liebig University Giessen, Germany, where they had been cultured for several years before the experiment (Tab. S4). To increase the number of replicate colonies in Experiment 1 & 3, we added four and six colonies respectively of *P. verrucosa* and *S. pistillata* that were collected from the central Red Sea in March 2019 (Tab. S4). Colonies were cut into 3–5 cm fragments with an angle grinder (Multitool 3000-15, Dremel, The Netherlands) and suspended on nylon strings (Experiment 1 & 3) or mounted on self-made concrete bases (Experiment 2). Fragments were allowed to heal for four to nine weeks before transfer into the experimental system and start of a two-week acclimation period (i.e., before addition of microplastic).

### Tank setup and water parameters

All experiments were performed in 40-L glass tanks that were part of a 2,300 L artificial seawater recirculating system, containing other scleractinian corals and associated organisms in separate tanks. The outflows of the tanks were equipped with filters (mesh size: 65 µm) to retain particles within the tank (Fig. S3, S5). Inflowing water was additionally filtered to remove potential contamination from the recirculation system. Seawater was treated in a central technical tank to remove organic refuse with a protein skimmer (ATI PowerCone 250i, ATI-Aquaristik, Germany), to adsorb phosphate with a silicate filter (ATI-Aquaristik, Germany) filled with iron oxide granulate (AquaLight PHOS, fein 0.5–2 mm, Aqualight GmbH, Germany), and to reduce microbial load with a UVC sterilizer (RWUVC/20/1000, RuWal aquatech, Italy). An algal refugium (*Chaetomorpha* sp.) with reverse light cycle buffered pH levels to 8–8.2.

Snails (*Turbo* sp. and *Euplica* sp.) were added to each experimental tank to reduce algal growth. Each experimental tank was equipped with a ‘turnover’ pump (Submarine Water Pump, Resun S-700, Resun, China) and a current pump (easyStream pro ES-28, Aqualight GmbH, Bramsche/Lappenstuhl, Germany) that created a water current of 3.8 cm s^-1^ at the corals’ position. To submerge microplastic particles accumulating at the water surface, the outflow piece of the turnover pump ended approximately 1 cm above the water surface (Fig. S3, S5). Water exchange rate in the tanks was approx. 78 L h^-1^, equivalent of two full water exchanges per hour. Water temperature was feedback-controlled (GHL Temp Sensor digital, ProfiLux 3 and ProfiLux 4, GHL Advanced Technology GmbH, Germany) and held constant before the heat phase between 26 and 27 °C with submersible titanium heaters (Schego Heater 300 W, Schemel & Goetz GmbH, Germany). Salinity (Instant Ocean, Aquarium Systems, France) was monitored daily and kept between 34–36. Light was provided with white and blue LED spots (SunaECO LED, Tropical Marine Centre, UK) at 120 µmol photons m^-2^ s^-1^ (Experiment 1 & 3) and T5 white and blue lamps (ATI Aquaristik, Germany) at 180-200 µmol photons m^-2^ s^-1^ (Experiment 2) at a 10:14 light:dark cycle. Alkalinity (1.42 mmol L^-1^) and calcium (Ca^2+^: 390–400 mg L^-1^) were monitored online (Alkatronic, Focustronic, NL) and maintained with a calcium reactor (PF 1001, Deltec, Germany). Nutrient concentrations were manually checked once a week: PO_4_^3^-: <0.02 mg L^-1^ (Spectroquant, Merck, Germany), NO_3_^-^: <0.02 mg L^-1^ and NO_2_^-^: <0.01 mg L^-1^ (MQuantTM, Merck, Germany), and Mg^2+^: 1300 mg L^-1^ (Mg Profi Test, Salifert, The Netherlands). Daily experimental maintenance included cleaning of in- and outflow filters, resuspension of particles attached to the tank walls, and feeding of corals. For feeding, approx. 0.53 g of frozen copepods (Calanoide Copepoden, Zooschatz, Germany) were added to each tank together with diluted amino acid mixture (Pohl’s Xtra special, Korallenzucht.de Vertriebs GmbH, Germany).

### Photosynthetic efficiency

We measured the photosynthetic efficiency of microalgal symbionts in the coral fragments as a measure of symbiont health. Photosynthetic efficiency is known to be negatively affected during heat stress (Jones et al., 1998), while microplastic exposure seems to have a positive, maybe compensatory, effect on photosynthetic efficiency (Reichert et al., 2019). For this, we used a Pulse-Amplitude-Modulated fluorometer (PAM-2500, Walz, Germany) with a 6 mm diameter fiber optic cable fitted with a translucent tube to keep a stable distance to the coral surface at a 45° angle. Effective photosynthetic efficiency (ΔF/Fm’) was measured for each fragment at different positions (three replicate positions in experiment 1 & 2, two replicate positions in experiment 3; whenever possible given size or health status of the coral fragment) during daytime. Maximum photochemical efficiency (Fv/Fm) was measured once per fragment after 40 min of dark acclimation when lights turned off at the end of the day. In the experiments, photosynthetic efficiency was assessed before the addition of microplastic particles, during the microplastic exposure, and during the combined microplastic and heat exposure (Tab. S2).

### Assessment of coral bleaching

To assess coral bleaching during combined microplastic and heat stress, we took standardized pictures of the fragments at several time points during the experiment (Tab. S3). For this, coral fragments were documented with a digital SLR camera (Nikon D7000) in an evenly illuminated macro photo studio (80 x 80 x 80 cm, Life of Photo). Fragments were always positioned the same way between time points with the upper side facing the camera. To prepare the analysis of coral tissue brightness over time, the background color of each picture was removed using Adobe Photoshop CC 2014 software. Pictures of the fragments were then analyzed with a self-written Python script, which created color histograms of tissue brightness (red, blue, green, and gray channels). Mean brightness of each channel was calculated from histogram data and used for further analyses. Bleaching severity was calculated based on tissue brightness on a scale of 0–255, in which 0 corresponds to the color black and 255 corresponds to the color white. That is, the higher the value, the more bleached the coral fragment. Because the analysis of all channels resulted in the same patterns, we present data from the gray channel here.

### 3D-scanning and analysis of tissue necrosis

To study coral tissue necrosis, coral fragments were documented using 3D-scanning at the end of the experiments. For this, a handheld 3D scanner (Artec Spider 3D, Artec 3D, Luxembourg) was used together with scanning software Artec Studio 12 (Artec 3D, Luxembourg) according to previous studies (Reichert et al., 2016). Fragments were placed on a rotating plate and scans were captured within 60–90 seconds from two angles in two full rotations. Models were calculated with a final mesh size of 0.2 mm, using the following settings: fine serial registration, global registration (minimal distance: 10, iterations: 2000, based on texture and geometry), and sharp fusion (resolution 0.2 mm, fill holes by radius, max. hole radius: 5). Outlier removal settings were adjusted to the underlying scan quality. For tissue surface area determination, meshes were trimmed manually at the tissue border. To determine the degree of necrosis, all bleached and necrotic tissue was trimmed in a second mesh to assess only the healthy, photosynthetically active tissue surface area and expressed as percent of the total tissue surface area. All meshes were exported as Wavefront “.obj” files to MeshLab Visual Computing Lab-ISTI-CNR (v1.3.4 BETA, 2014) and the surface area was calculated using the “compute geometric measures” tool.

### Survival analysis

In the first experiment, we assessed coral mortality in response to combined impact of heat stress and different concentrations of microplastic. For this, each fragment was visually checked twice a day and recorded as dead when no tissue was left on the skeleton.

### Statistical analyses

All statistical analyses and data plots were performed in the R statistical environment (v.3.6.1 (R Core Team, 2019)). The effect of (i) heat stress and microplastic exposure, (ii) microplastic in the initial phase of the experiments (only in Experiments 1 and 2), and of (iii) microplastic under heat stress in the following phase of the experiment, on photosynthesis (ΔF/F’m and Fv/Fm), tissues brightness (gray scale) and necrosis (percent healthy tissue) were analyzed using linear mixed-effect (lme) models. Individual models were constructed for each response variable and species. Microplastic treatment, heat stress and its interaction (i) or the microplastic treatments alone (ii and iii) were selected as fixed effects. In experiment 1, this impact was analyzed based on all microplastic treatments pooled as well as on the four concentration levels individually. As random effects, colony, origin (Indopacific or Red Sea), and fragment identity were included in the model if applicable. Data were transformed (squared) if necessary, prior to analysis to ensure that test assumptions were met. Residual structures were checked visually by graphical residual analysis. The models were fitted using the lme4 package v.1.1-14 (Bates et al., 2015). Detailed specifications on the individual model equations are provided in Tab. S5, 6, and 7. The effect of the microplastic treatments (individual and pooled concentrations) on coral mortality was analyzed using mixed-effect Cox models, based on cumulative incidence analyzes (Cox, 1972; Ripatti and Palmgren, 2000; Therneau et al., 2003). The microplastic treatment was included as fixed effect, and colony and origin as random effects. The effect of heat stress exposure on coral mortality was analyzed using log-rank statistics, based on a two-Sample tests for censored data (Hothorn et al., 2008), as no events occurred in the non-heated group, which resulted in degenerated estimates when using mixed-effect Cox models. One model was constructed for each species. The models were fitted using the coxme package v. 2.2-16 and the coin package 1.3-1, and plots were generated using the survival package v.2.41-3 (Hothorn et al., 2008; Therneau, 2020, 1999). To compare the magnitude of the effects of heat stress and microplastic exposure on coral holobiont health, we visualized the z-value statistics for each lme model parameter in a heatmap. Data visualization was performed using the R package ggplot2 (Wickham, 2016).

## Results

### Effect of different concentrations of microplastic on coral heat stress tolerance

The cumulative effects of PE microplastic (i.e., in different concentrations) and heat stress (i.e., whether microplastic aggravates the effects of heat stress) on the two reef-building corals species *Pocillopora verrucosa* and *Stylophora pistillata* were studied in the first experiment. Maximum photosynthetic efficiency (Fv/Fm) of the microalgal symbionts significantly decreased during the heat stress treatment of microplastic-free control and microplastic treated fragments of both coral species (*P. verrucosa*, z = −9.51, p < 0.0001; *S. pistillata*, z = −13.559, p < 0.0001; Fig. 2a, Fig. S11, Tab. S5). In *P. verrucosa*, Fv/Fm was not significantly different between control and microplastic treatments regardless of heat stress (p > 0.05). In *S. pistillata*, Fv/Fm of control and microplastic treatment was similar at ambient temperature (p > 0.05), with significantly higher values in the highest microplastic treatment compared to the control under heat stress (2 500 particles L^-1^, z = − 3.136, p = 0.00686). Effective photosynthetic efficiency (ΔF/Fm’) of the microalgal symbionts was also significantly reduced in response to heat stress (*P. verrucosa*, z = −16.182, p < 0.0001; *S. pistillata*, z = −17.25, p < 0.0001; Fig. S12, Tab. S5). In *P. verrucosa*, ΔF/Fm’ was significantly increased in the highest microplastic treatment compared to the control under heat stress (2 500 particles L^-1^, z = −2.863, p = 0.0168), but not at ambient temperatures (p > 0.05). In *S. pistillata*, ΔF/Fm’ was significantly increased in the highest microplastic treatment compared to the control under ambient temperature (2 500 particles L^-1^, z = −2.508, p = 0.0485) and under heat stress (2 500 particles L^-1^, z = −3.541, p = 0.0016).

**Figure 2.**
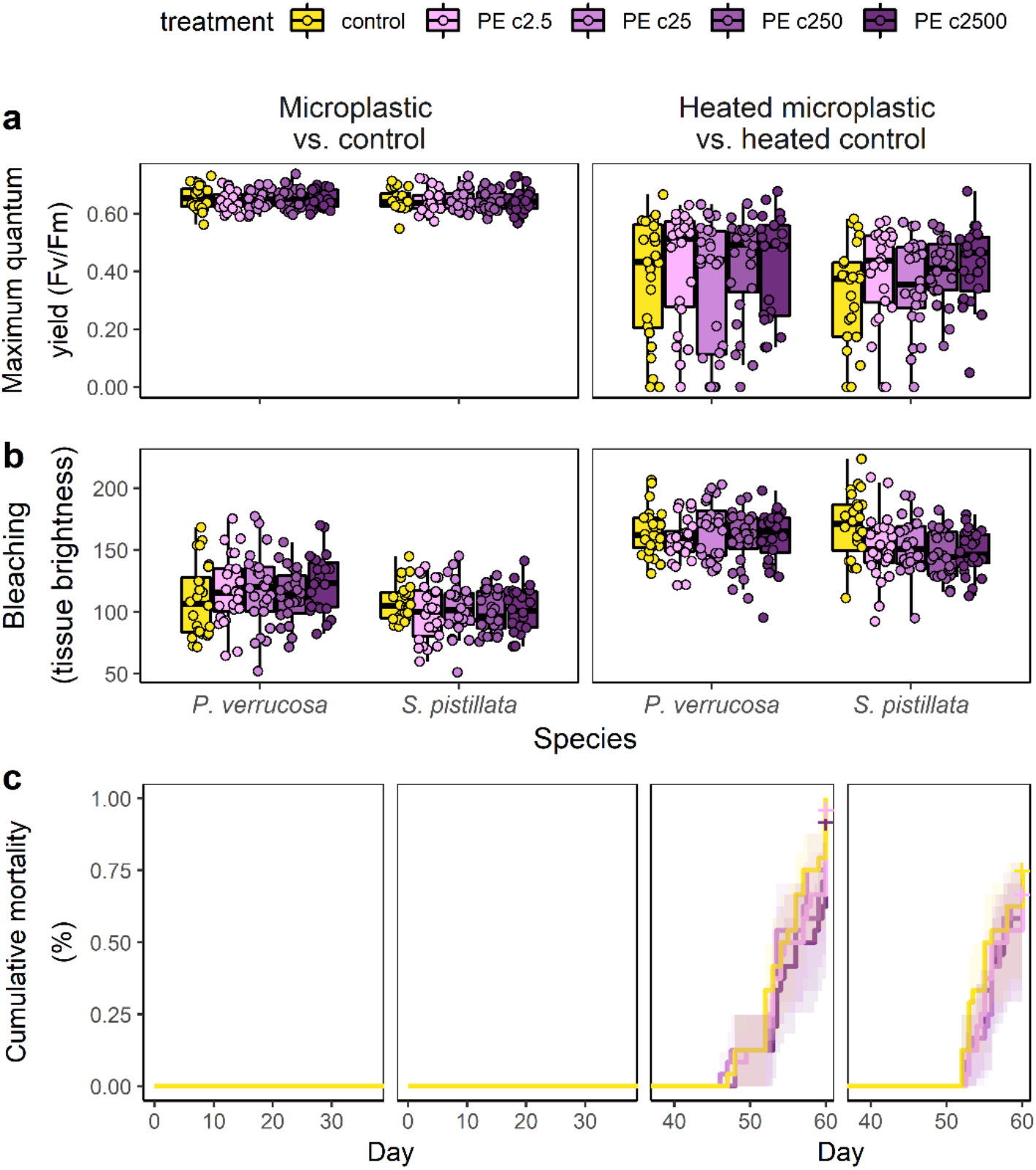
Effect of different microplastic particle concentrations in combination with heat stress on corals. Final time points from the time series of the combined PE-microplastic and heat stress experiment on two stony coral species (*Pocillopora verrucosa, Stylophora pistillata)*. In both species, coral performance, assessed as a) maximum photosynthetic efficiency of algal symbionts, b) bleaching, and c) cumulative mortality, was mainly affected by heat stress. Microplastic treated corals at different PE particle concentrations (2.5, 25, 250, and 2 500 particles L^-1^) showed only a small microplastic treatment response in comparison to microplastic-free controls. Bleaching displayed as tissue brightness ranging from 0 = black to 255 = white.

Both coral species bleached severely in response to the heat stress treatment (*P. verrucosa*, z = 9.229, p < 0.0001; *S. pistillata*, z = −11.338, p < 0.0001; Fig. 2b, Fig. S13, Tab. S5), but not due to exposure to microplastic particles alone, even at very high concentrations (all p > 0.05, Fig. S13– 21). In the heat stress treatment, bleaching severity was similar between microplastic-free control and different concentrations of particles in *P. verrucosa* (p > 0.05). In *S. pistillata*, microplastic treated corals bleached less under heat stress than their microplastic free controls (all pairwise comparisons, z = 2.682-3.852, p = 0.015-0.0005, respectively).

Both coral species experienced high mortality when subjected to heat stress (Fig. 2c). At ambient temperature, no mortality was observed in microplastic treatment and control of either species (p > 0.05), with significantly increased mortality rates after the onset of the heat stress (*P. verrucosa*, z = 14.702, p < 0.0001; *S. pistillata*, z = 11.248, p < 0.0001). Microplastic treatment had no effect on mortality rates under heat stress in *P. verrucosa* (p > 0.05). In contrast, under heat stress the control treatment in *S. pistillata* experienced significantly higher mortality compared to all concentrations of microplastic (z = −3.83, p = 0.0001).

### Effect of microplastic on heat tolerance of different stony coral species

The combined effect of microplastic pollution (PE particles at 230 particles L^-1^) and heat stress was tested on five coral species representing different life history strategies, growth forms, and environmental susceptibilities (i.e., *Acropora muricata, Montipora digitata, Porites cylindrica, P. verrucosa*, and *S. pistillata*) was tested in the second experiment. In congruence with the previous experiment, the effect of heat stress was consistently more severe than the effect of microplastic exposure on all coral species (Fig. 3). Maximum photosynthetic efficiency (Fv/Fm) of the microalgal symbionts significantly decreased in control and microplastic treatments of all coral species when subjected to heat (Fig. 3a, Tab. S6). This trend started at the beginning of the heat period and continued to decrease with more severe and protracted heat stress (Fig. S22). *P. verrucosa* was the only species with different Fv/Fm between control and MP treatment, with significantly reduced values in the microplastic treatment compared to the control at ambient temperatures (z = −2.733, p = 0.0063) and under heat stress (z = −2.086, p = 0.037). Effective photosynthetic efficiency (ΔF/Fm’) of the microalgal symbionts was affected less by the treatments, with significant reductions in response to heat stress only for *A. muricata* (z = −4.022, p < 0.0001) and *P. verrucosa* (z = −4.653, p < 0.0001) (Fig. S23, Tab. S6). MP treatment had no significant effect on ΔF/Fm’ compared to microplastic-free controls (p > 0.05).

**Figure 3.**
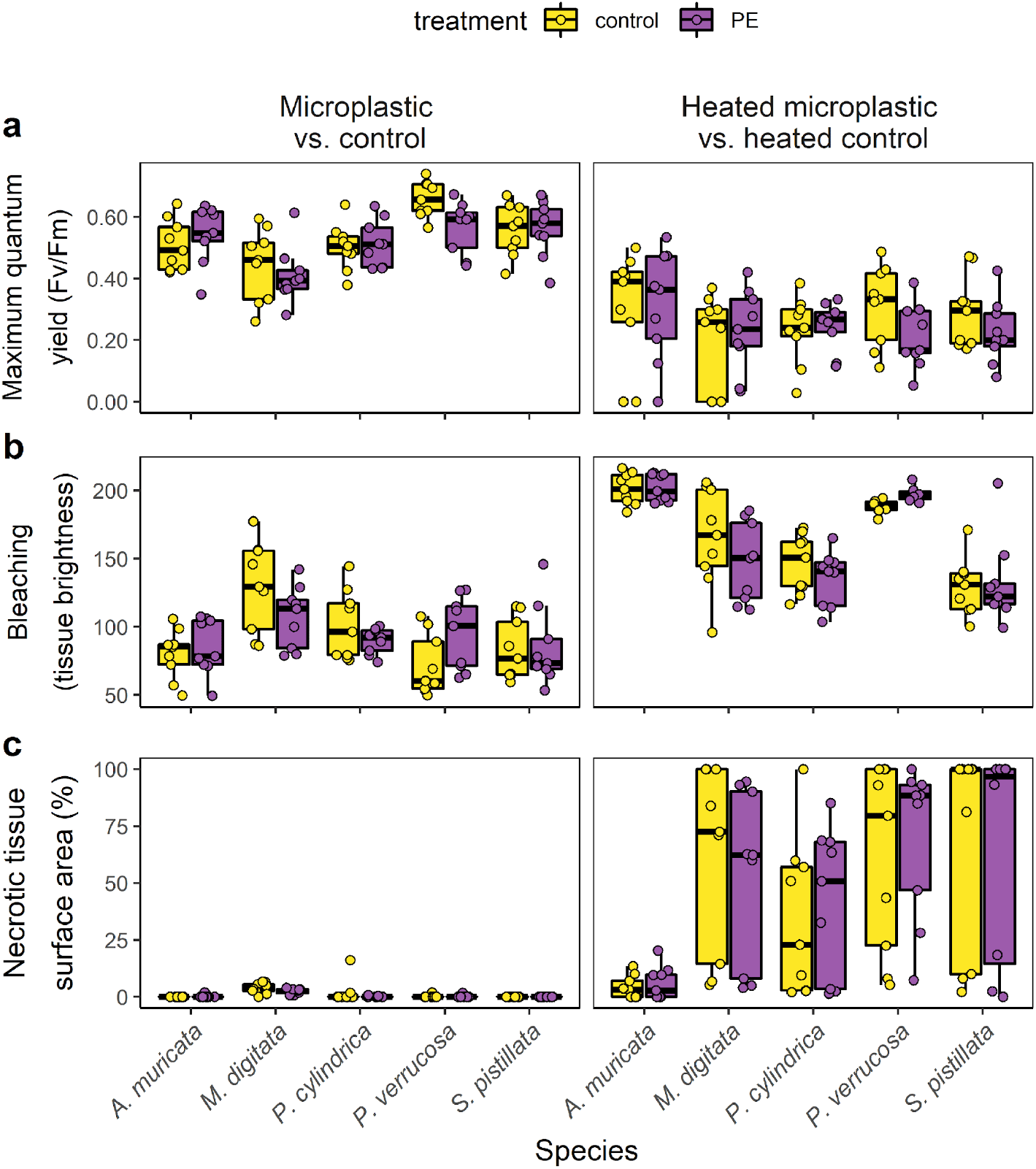
Effect of combined microplastic treatment and heat stress on different coral genera. Final time points from the time series of combined PE-microplastic and heat stress experiment on five stony coral species (*Acropora muricata, Montipora digitata, Porites cylindrica, Pocillopora verrucosa, Stylophora pistillata*). Coral performance, assessed as a) maximum photosynthetic efficiency of algal symbionts, b) bleaching, and c) necrosis, was mainly affected by heat stress for all species. In comparison, microplastic-free controls and microplastic treated corals showed only small differences in performance. Bleaching displayed as tissue brightness ranging from 0 = black to 255 = white.

All coral species significantly bleached in response to the heat stress treatment (Fig. 3b, all p < 0.001, Fig. S24–29, Tab. S6). Microplastic exposure had comparatively small, but species-specific effects on coral heat tolerance. Bleaching severity was significantly different between microplastic treatment and microplastic-free control in *M. digitata* and *P. verrucosa*, while *A. muricata, P. cylindrica*, and *S. pistillata* were not affected (Fig. S24). Whereas under heat stress microplastic treated corals of *M. digitata* bleached less than microplastic-free controls (z = −2.277, p = 0.0228), microplastic treated corals of *P. verrucosa* were more bleached (i.e., paler) than controls under ambient temperatures (z = 2.544, p = 0.0109) and heat stress (z = 2.821, p = 0.0048). All coral species showed significant tissue necrosis in response to the heat stress at the end of the experiment (Fig. 3c). The proportion of necrotic tissue between control and microplastic treatment was not significantly different (p > 0.05).

### Effect of different mixtures of microplastic on coral heat stress tolerance

The combined effect of different mixtures of microplastic particles (i.e., fibers, tirewear, beach, PE) and heat stress on two reef-building coral species were tested in the third experiment. Different mixtures of microplastic particles had minor effects on the heat stress tolerance of *P. verrucosa* compared to microplastic-free heated controls, but no detectable effect on *S. pistillata* (Fig. 4). Maximum photosynthetic efficiency (Fv/Fm) of the microalgal symbionts was similar between heated microplastic and heated control treatments of both species (p > 0.05; Fig. 4a, Tab. S7). Effective quantum yield (ΔF/Fm’) accordingly showed no strong response between heated microplastic treated and heated control corals (p > 0.05; Fig. S30, Tab. S7). The only exception was a significant reduction of 20 % in ΔF/Fm’ of *P. verrucosa* in the fibers treatment (z = 3.176, p = 0.006). Both species bleached and underwent tissue necrosis in response to the heat stress treatment (Fig. S31, 32), however the degree of bleaching and necrosis was not significantly different between microplastic-exposed and control corals (p > 0.05, Fig. 4b, c, Tab. S7).

**Figure 4.**
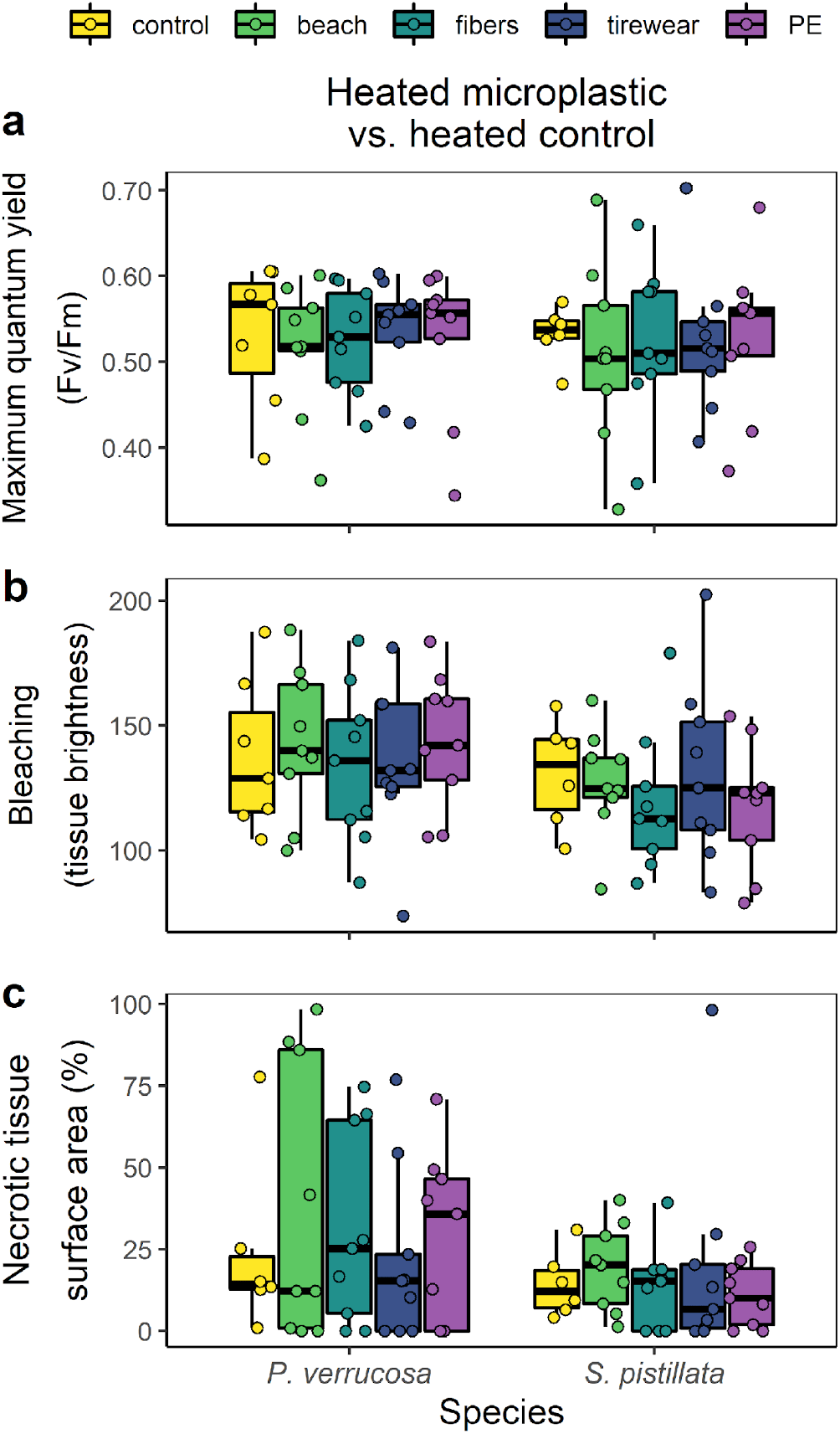
Effect of different types of microplastic in combination with heat stress on corals. Summary of main results at the end of combined microplastic and heat stress experiment on two stony coral species (*Pocillopora verrucosa, Stylophora pistillata)*. Microplastic-free controls and microplastic-treated corals showed a strong heat stress response in performance parameters a) maximum photosynthetic efficiency of algal symbionts, b) bleaching, and c) tissue necrosis. In comparison, microplastic treatments of different compositions (secondary beach plastic, artificial clothing fibers, tirewear from the automobile sector, and PE particles) had no significant effects on coral heat stress tolerance. Bleaching displayed as tissue brightness ranging from 0 = black to 255 = white.

### Comparison of effect sizes of microplastic and heat stress

In summary, all three experiments showed that microplastic exposure had no cumulative negative effect on corals’ thermal tolerance under heat stress, and differences between heated microplastic-free controls and heated microplastic-treated corals were minor (Fig. 2-4). In a comparative framework the effect of heat stress on coral health and performance was orders of magnitude stronger than the effect of microplastic exposure (Fig. 5). This pattern was consistent across three independent experiments, in which different microplastic types and concentrations and different protocols for heat stress exposure were used, testifying to a strong, reproducible effect.

**Figure 5.**
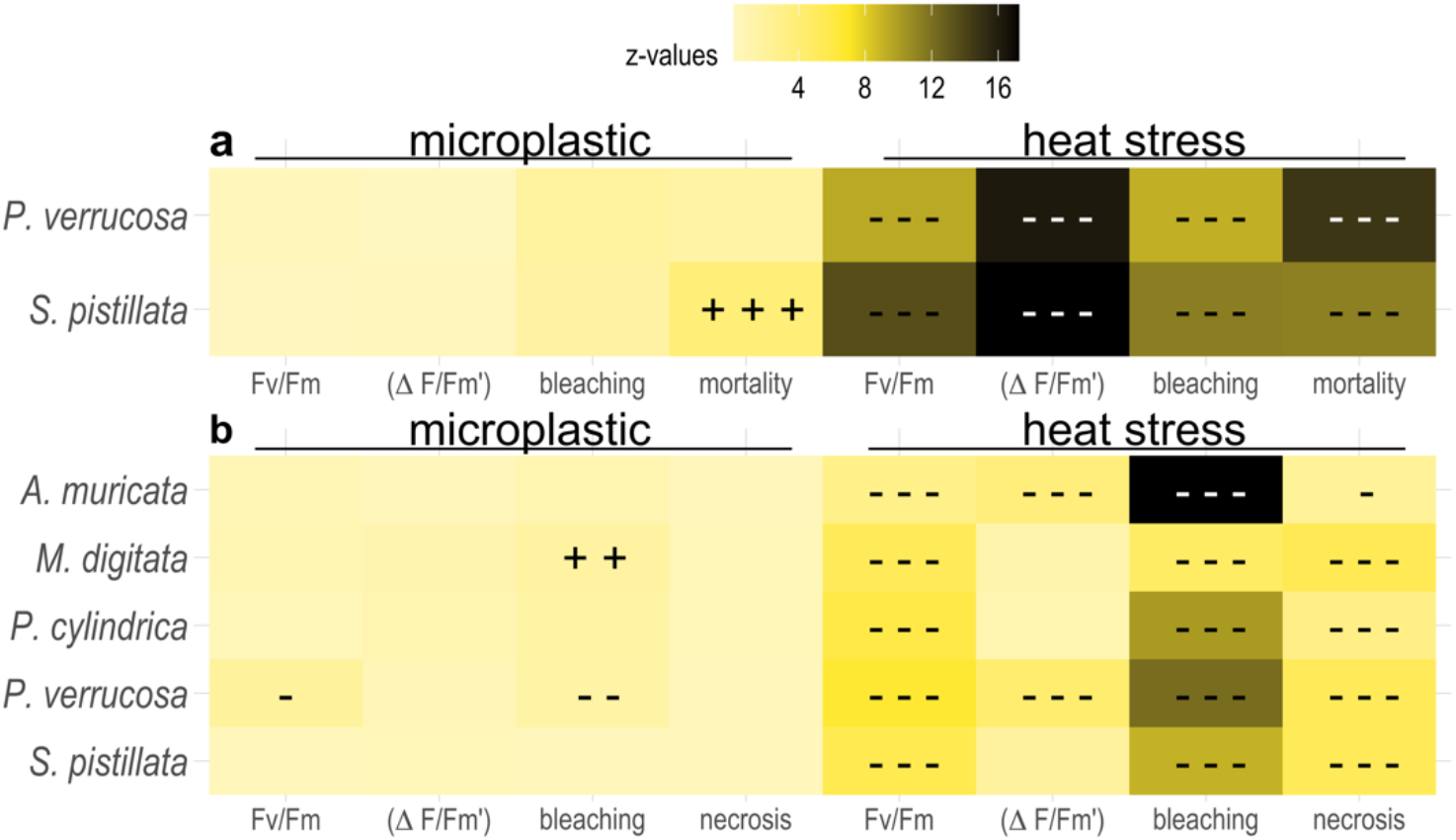
Comparison of effect sizes between microplastic and heat stress treatments. The effect of heat stress on coral health and performance is much stronger than the effect of microplastic treatment. The effect of heat stress and PE microplastic pollution were compared in independent experiments a) testing the effect of different particle concentrations on two stony coral species (*Pocillopora verrucosa, Stylophora pistillata*) and b) testing the species-specific responses of five stony coral species (*Acropora muricata, Montipora digitata, Porites cylindrica, P. verrucosa, S. pistillata*). Five response parameters were assessed as measure of health and performance: maximum quantum yield, effective quantum yield, bleaching, necrosis, and mortality. Strength of effect shown in heatmap based on z-values derived from linear mixed-effects models. - / -- / --- and + / ++ / +++ signify that the stressor (microplastic or heat) imparts a negative (-) or positive (+) effect compared to the control group at significance levels of p < 0.05 / 0.01 / 0.001, respectively.

## Discussion

Microplastic exposure had only a minor and infrequent cumulative negative effect on coral thermal tolerance under heat stress, regardless of particle concentrations (even beyond projected future concentrations) and particle types. The negligible effect of microplastic on coral thermal tolerance we found here, is in stark contrast to other additive stressors such as eutrophication (Wiedenmann et al., 2013), which increase the susceptibility of corals to bleaching. This pattern appeared to be consistent across a broad range of reef-building coral genera. The only exception was found for the exposure to fibers, which negatively impacted the photophysiology of *P. verrucosa* and increased the severity of heat stress. Microplastic in the form of fibers might cause more stress than particles, as fibers tend to get entangled on the surface of coral colonies (Hierl et al., 2021) and decrease photosynthetic efficiency of algal symbionts at ambient temperatures (Mendrik et al., 2021). This is also evidenced by higher deposition rates of fibers into coral skeletons compared to particles (Krishnakumar et al., 2021). Notably, microplastic exposure had small but surprisingly beneficial effects on the heat tolerance of reef-building corals in some cases (e.g., in *S. pistillata* during Experiment 1 and in *M. digitata* during Experiment 2). These may be related to increased photosynthetic efficiency of the algal symbionts, a potentially compensatory effect that was previously described in response to microplastic exposure (Lanctôt et al., 2020; Reichert et al., 2019). Furthermore, microplastic exposure has been shown to induce a response at the molecular level by upregulating the expression of heat shock proteins (Corinaldesi et al., 2021). This is also a common response in corals exposed to temperature stress (e.g., Rosic et al., 2011; Seveso et al., 2014). Therefore, the more permanent microplastic exposure may prepare the corals for additional short-term (heat) stress through the process of molecular frontloading (Barshis et al., 2013). Although the impact of microplastic on the feeding behavior of corals is not well understood yet (Allen et al., 2017; Axworthy and Padilla-Gamiño, 2019; Chapron et al., 2018), another potential explanation for better performance in the microplastic treatments could be higher heterotrophic feeding rates stimulated by the presence of microplastic particles. Higher energy reserves and heterotrophic capacity may lead to higher thermal tolerance and shorter recovery times after coral bleaching (Grottoli et al., 2006). Thus, corals might indirectly benefit from microplastic exposure in the short term, which might help explain the lack of an additive effect. However, microplastic pollution is a constant stressor in coral reef ecosystems. Over longer time scales, such compensatory effects might not be sustainable, requiring further confirmation.

Microplastic alone had negligible, sometimes beneficial effects on the physiology of the corals, for common microplastic scenarios up to 250 particles L^-1^. Only the highest concentration (i.e., 2,500 particles L^-1^), which was chosen as a high pollution scenario, had detectable impacts on the physiology of the tested species and caused both negative (decreased photosynthetic efficiency) and positive (lower bleaching and mortality) effects. This is in line with previous studies showing that microplastics have only small effects when tested for common pollution scenarios (Berry et al., 2019; Grillo et al., 2021). Even though the rough surface and complex shape of coral colonies increase the chances to get in contact with microplastics in the environment (de Smit et al., 2021; Lim et al., 2020), at low environmental concentrations these coincidences are probably infrequent and largely stochastic. Under (unrealistically) high concentrations, corals have been shown to respond to microplastics on various organismal levels, including, photophysiology, immunity, growth, and health (Lanctôt et al., 2020; Tang et al., 2018). Changes caused by high particle concentrations might reflect an early stage of the disruption of the coral-algae symbiosis (Okubo et al., 2018; Su et al., 2020). While studies exposing corals to unrealistically high concentrations help to explain the potential impact and pathways in which microplastics may affect coral health, artificially exposing corals to ‘microplastic storms’ may be of little relevance to corals in the reef environment. Thus, particle concentrations applied to test the ecological relevance of microplastics on corals should be guided by environmental concentrations to better relate the true impacts of the stressor.

In accordance with previous studies our findings also show that the effects of microplastic exposure alone are species-specific, with *P. verrucosa* appearing to be more likely impacted by microplastic pollution than other coral species (Mouchi et al., 2019; Reichert et al., 2019). Similarly, *Acropora* sp. was more susceptible to microplastics than *Seriatopora hystrix* in a comparative study (Mendrik et al., 2021). When more species were tested together, *P. verrucosa* and *A. muricata* have been shown to suffer from mortality and reduced growth compared to *P. lutea* and *Heliopora coerulea*, which were largely unaffected (Reichert et al., 2019). What unites *P. verrucosa* and *Acropora* sp. is their more frequent reaction to microplastics compared to other species (Hall et al., 2015; Reichert et al., 2018). Reasons for this might be their high morphological complexity (Reichert et al., 2017), which confers specific boundary layer characteristics to coral colonies and increases chances of contact to microplastics (Lim et al., 2020). Also differences in feeding mode of the tested coral species (e.g., active capture feeding in *A. muricata* and *P. verrucosa* or feeding on dissolved organic matter in *P. lutea*), as well as cleaning efficiency might confer species-specific the differences in microplastic handling (Houlbrèque and Ferrier-Pagès, 2009). The more frequent handling of the particles may be energetically costly, which in turn may lead to higher bleaching susceptibility.

## Conclusions

The results of this study not only show that microplastics can have both aggravating and mitigating impacts on corals under heat stress, but highlight that the effects of microplastics appear small when compared to the impacts of heat. Further investigations are needed to analyze pathways of adverse and beneficial effects microplastics have on corals, addressing molecular processes as well as the impacts of plastic-associated microbiota or toxins under ocean warming scenarios. The rare occurrence of additive effects on the susceptibility of corals to bleaching underlines the ambivalence of this stressor. These insights have strong implications for setting priorities in individual action, local and regional conservation and coastal management practices, and global policy. While efforts to reduce plastic pollution should continue, they should not replace more urgent efforts to cut global carbon dioxide emissions, which are immediately needed to preserve remaining coral reef ecosystems.

## Acknowledgements

We thank Susanne Kühn (Wageningen University and Research, The Netherlands) for supply with beach microplastic and Sebastian Primpke (Alfred-Wegener-Institut, Helgoland, Germany) for the FTIR characterization of the fiber treatment in experiment 3. We thank Olivia Metz (Justus Liebig University Giessen, Germany) for support during Experiment 2, and Birgit Zettl and Christina Anding (both Justus Liebig University Giessen, Germany) for animal care and maintenance. This study was conducted as part of the ‘Ocean2100’ global change simulation project of the Colombian-German Center of Excellence in Marine Sciences (CEMarin) funded by the German Academic Exchange Service.

## Author Contributions

JR, MZ conceived the study; JR, VT, RA, KB, JK, PS, MZ performed experiments; JR analyzed data; JR, MZ, TW interpreted data; TW acquired funding; JR, MZ wrote the manuscript with contributions from VT, TW. All authors read and approved the final manuscript.

## Data and materials availability

All raw data for this study is provided with the published article.

## References

Ainsworth, T.D., Heron, S.F., Ortiz, J.C., Mumby, P.J., Grech, A., Ogawa, D., Eakin, C.M., Leggat, W., 2016. Climate change disables coral bleaching protection on the Great Barrier Reef. Science 352, 338–342. doi:10.1126/science.aac7125

Allen, A.S., Seymour, A.C., Rittschof, D., 2017. Chemoreception drives plastic consumption in a hard coral. Mar. Pollut. Bull. 124, 198–205. doi:10.1016/j.marpolbul.2017.07.030

Andrady, A.L., 2017. The plastic in microplastics: A review. Mar. Pollut. Bull. 119, 12–22. doi:10.1016/j.marpolbul.2017.01.082

Axworthy, J.B., Padilla-Gamiño, J.L., 2019. Microplastics ingestion and heterotrophy in thermally stressed corals. Sci. Rep. 9, 1–8. doi:10.1038/s41598-019-54698-7

Badylak, S., Phlips, E., Batich, C., Jackson, M., Wachnicka, A., 2021. Polystyrene microplastic contamination versus microplankton abundances in two lagoons of the Florida Keys. Sci. Rep. 11, 1–10. doi:10.1038/s41598-021-85388-y

Barshis, D.J., Ladner, J.T., Oliver, T.A., Seneca, F.O., Traylor-Knowles, N., Palumbi, S.R., 2013. Genomic basis for coral resilience to climate change. Proc. Natl. Acad. Sci. U. S. A. 110, 1387– 1392. doi:10.1073/pnas.1210224110

Bates, D., Mächler, M., Bolker, B.M., Walker, S.C., Machler, M., Bolker, B.M., Walker, S.C., 2015. Fitting Linear Mixed-Effects Models Using lme4. J. Stat. Softw. 67, 1–48. doi:10.18637/jss.v067.i01

Berry, K.L.E., Epstein, H.E., Lewis, P.J., Hall, N.M., Negri, A.P., 2019. Microplastic Contamination Has Limited Effects on Coral Fertilisation and Larvae. Diversity 11, 228. doi:10.3390/d11120228

Browne, M.A., Crump, P., Niven, S.J., Teuten, E., Tonkin, A., Galloway, T., Thompson, R., 2011. Accumulation of microplastic on shorelines woldwide: Sources and sinks. Environ. Sci. Technol. 45, 9175–9179. doi:10.1021/es201811s

Carr, S.A., 2017. Sources and dispersive modes of micro-fibers in the environment. Integr. Environ. Assess. Manag. 13, 466–469. doi:10.1002/ieam.1916

Chapron, L., Peru, E., Engler, A., Ghiglione, J.F., Meistertzheim, A.L., Pruski, A.M., Purser, A., Vétion, G., Galand, P.E., Lartaud, F., 2018. Macro- and microplastics affect cold-water corals growth, feeding and behaviour. Sci. Rep. 8, 15299. doi:10.1038/s41598-018-33683-6

Corinaldesi, C., Canensi, S., Dell’Anno, A., Tangherlini, M., Di Capua, I., Varrella, S., Willis, T.J., Cerrano, C., Danovaro, R., 2021. Multiple impacts of microplastics can threaten marine habitat-forming species. Commun. Biol. 4, 431. doi:10.1038/s42003-021-01961-1

Corona, E., Martin, C., Marasco, R., Duarte, C.M., 2020. Passive and active removal of marine microplastics by a mushroom coral (Danafungia scruposa). Front. Mar. Sci. 7, 1–9. doi:10.3389/fmars.2020.00128

Cox, D.R., 1972. Regression models and life-tables. J. R. Stat. Soc. Ser. B 34, 187–220. doi:10.1007/978-1-4612-4380-9_37

de Smit, J.C., Anton, A., Martin, C., Rossbach, S., Bouma, T.J., Duarte, C.M., 2021. Habitat-forming species trap microplastics into coastal sediment sinks. Sci. Total Environ. 772, 145520. doi:10.1016/j.scitotenv.2021.145520

Fisher, R., O’Leary, R.A., Low-Choy, S., Mengersen, K., Knowlton, N., Brainard, R.E., Caley, M.J., 2015. Species richness on coral reefs and the pursuit of convergent global estimates. Curr. Biol. 25, 500–505. doi:10.1016/j.cub.2014.12.022

Franco, A., Rückert, C., Blom, J., Busche, T., Reichert, J., Schubert, P., Goesmann, A., Kalinowski, J., Wilke, T., Kämpfer, P., Glaeser, S.P., 2020. High diversity of Vibrio spp. associated with different ecological niches in a marine aquaria system and description of Vibrio aquimaris sp. nov. Syst. Appl. Microbiol. 43, 126123. doi:10.1016/j.syapm.2020.126123

Frölicher, T.L., Fischer, E.M., Gruber, N., 2018. Marine heatwaves under global warming. Nature 560, 360–364. doi:10.1038/s41586-018-0383-9

Grillo, J.F., Sabino, M.A., Ramos, R., 2021. Short-term ingestion and tissue incorporation of Polystyrene microplastic in the scleractinian coral Porites porites. Reg. Stud. Mar. Sci. 43, 101697. doi:10.1016/j.rsma.2021.101697

Grottoli, A.G., Rodrigues, L.J., Palardy, J.E., 2006. Heterotrophic plasticity and resilience in bleached corals. Nature 440, 1186–9. doi:10.1038/nature04565

Hall, N.M., Berry, K.L.E., Rintoul, L., Hoogenboom, M.O., 2015. Microplastic ingestion by scleractinian corals. Mar. Biol. 162, 725–732. doi:10.1007/s00227-015-2619-7

Hartmann, N.B., Hüffer, T., Thompson, R.C., Hassellöv, M., Verschoor, A., Daugaard, A.E., Rist, S., Karlsson, T., Brennholt, N., Cole, M., Herrling, M.P., Hess, M.C., Ivleva, N.P., Lusher, A.L., Wagner, M., 2019. Are we speaking the same language? Recommendations for a definition and categorization framework for plastic debris. Environ. Sci. Technol. 53, 1039–1047. doi:10.1021/acs.est.8b05297

Hierl, F., Wu, H.C., Westphal, H., 2021. Scleractinian corals incorporate microplastic particles: identification from a laboratory study. Environ. Sci. Pollut. Res. doi:10.1007/s11356-021-13240-x

Hoegh-Guldberg, O., 1999. Climate change, coral bleaching and the future of the world’s coral reefs. Mar. Freshw. Res. 50, 893–66. doi:10.1071/MF99078

Hothorn, T., Bretz, F., Westfall, P., 2008. Simultaneous inference in general parametric models. Biometrical J. 50, 346–363. doi:10.1002/bimj.200810425

Houlbrèque, F., Ferrier-Pagès, C., 2009. Heterotrophy in tropical scleractinian corals. Biol. Rev. 84, 1– 17. doi:10.1111/j.1469-185X.2008.00058.x

Hughes, T.P., Anderson, K.D., Connolly, S.R., Heron, S.F., Kerry, J.T., Lough, J.M., Baird, A.H., Baum, J.K., Berumen, M.L., Bridge, T.C., Claar, D.C., Eakin, C.M., Gilmour, J.P., Graham, N.A.J., Harrison, H., Hobbs, J.-P.A., Hoey, A.S., Hoogenboom, M., Lowe, R.J., McCulloch, M.T., Pandolfi, J.M., Pratchett, M., Schoepf, V., Torda, G., Wilson, S.K., 2018. Spatial and temporal patterns of mass bleaching of corals in the Anthropocene. Science 359, 80–83. doi:10.1126/science.aan8048

Jensen, L.H., Motti, C.A., Garm, A.L., Tonin, H., Kroon, F.J., 2019. Sources, distribution and fate of microfibres on the Great Barrier Reef, Australia. Sci. Rep. 9, 1–15. doi:10.1038/s41598-019-45340-7

Jones, R.J., Hoegh-Guldberg, O., Larkum, A.W.D., Schreiber, U., 1998. Temperature-induced bleaching of corals begins with impairment of the CO2 fixation mechanism in zooxanthellae. Plant, Cell Environ. 21, 1219–1230. doi:10.1046/j.1365-3040.1998.00345.x

Kirstein, I. V., Kirmizi, S., Wichels, A., Garin-Fernandez, A., Erler, R., Löder, M., Gerdts, G., 2016. Dangerous hitchhikers? Evidence for potentially pathogenic Vibrio spp. on microplastic particles. Mar. Environ. Res. 120, 1–8. doi:10.1016/j.marenvres.2016.07.004

Koelmans, A.A., Kooi, M., Law, K.L., van Sebille, E., 2017. All is not lost: deriving a top-down mass budget of plastic at sea. Environ. Res. Lett. 12, 114028. doi:10.1088/1748-9326/aa9500

Konkel, M.E., Tilly, K., 2000. Temperature-regulated expression of bacterial virulence genes. Microbes Infect. 2, 157–166. doi:10.1016/S1286-4579(00)00272-0

Krishnakumar, S., Anbalagan, S., Hussain, S.M., Bharani, R., Godson, P.S., Srinivasalu, S., 2021. Coral annual growth band impregnated microplastics (Porites sp.): a first investigation report. Wetl. Ecol. Manag. 9. doi:10.1007/s11273-021-09786-9

Kroon, F.J., Motti, C.E., Jensen, L.H., Berry, K.L.E., 2018. Classification of marine microdebris: A review and case study on fish from the Great Barrier Reef, Australia. Sci. Rep. 8, 1–15. doi:10.1038/s41598-018-34590-6

Kühn, S., van Oyen, A., Booth, A.M., Meijboom, A., van Franeker, J.A., 2018. Marine microplastic: Preparation of relevant test materials for laboratory assessment of ecosystem impacts. Chemosphere 213, 103–113. doi:10.1016/j.chemosphere.2018.09.032

Lamb, J.B., Willis, B.L., Fiorenza, E.A., Couch, C.S., Howard, R., Rader, D.N., True, J.D., Kelly, L.A., Ahmad, A., Jompa, J., Harvell, C.D., 2018. Plastic waste associated with disease on coral reefs. Science 359, 460–462. doi:10.1126/science.aar3320

Lanctôt, C.M., Bednarz, V.N., Melvin, S., Jacob, H., Oberhaensli, F., Swarzenski, P.W., Ferrier-Pagès, C., Carroll, A.R., Metian, M., 2020. Physiological stress response of the scleractinian coral Stylophora pistillata exposed to polyethylene microplastics. Environ. Pollut. 263, 114559. doi:10.1016/j.envpol.2020.114559

Lim, H.S., Fraser, A., Knights, A.M., 2020. Spatial arrangement of biogenic reefs alters boundary layer characteristics to increase risk of microplastic bioaccumulation. Environ. Res. Lett. 15. doi:10.1088/1748-9326/ab83ae

Martin, C., Corona, E., Mahadik, G.A., Duarte, C.M., 2019. Adhesion to coral surface as a potential sink for marine microplastics. Environ. Pollut. 255, 113281. doi:10.1016/j.envpol.2019.113281

Mendrik, F.M., Henry, T.B., Burdett, H., Hackney, C.R., Waller, C., Parsons, D.R., Hennige, S.J., 2021. Species-specific impact of microplastics on coral physiology. Environ. Pollut. 269, 116238. doi:10.1016/j.envpol.2020.116238

Montano, S., Seveso, D., Maggioni, D., Galli, P., Corsarini, S., Saliu, F., 2020. Spatial variability of phthalates contamination in the reef-building corals Porites lutea, Pocillopora verrucosa and Pavona varians. Mar. Pollut. Bull. 155, 111117. doi:10.1016/j.marpolbul.2020.111117

Mouchi, V., Chapron, L., Peru, E., Pruski, A.M., Meistertzheim, A.-L., Vétion, G., Galand, P.E., Lartaud, F., 2019. Long-term aquaria study suggests species-specific responses of two cold-water corals to macro-and microplastics exposure. Environ. Pollut. 253, 322–329. doi:10.1016/j.envpol.2019.07.024

Okubo, N., Takahashi, S., Nakano, Y., 2018. Microplastics disturb the anthozoan-algae symbiotic relationship. Mar. Pollut. Bull. 135, 83–89. doi:10.1016/j.marpolbul.2018.07.016

Patterson, J., Jeyasanta, K.I., Sathish, N., Edward, J.K.P., Booth, A.M., 2020. Microplastic and heavy metal distributions in an Indian coral reef ecosystem. Sci. Total Environ. 744, 140706. doi:10.1016/j.scitotenv.2020.140706

Primpke, S., Wirth, M., Lorenz, C., Gerdts, G., 2018. Reference database design for the automated analysis of microplastic samples based on Fourier transform infrared (FTIR) spectroscopy. Anal. Bioanal. Chem. 410, 5131–5141. doi:10.1007/s00216-018-1156-x

R Core Team, 2019. R: A language and environment for statistical computing [WWW Document]. URL http://www.r-project.org/

Reichert, J., Arnold, A.L., Hoogenboom, M.O., Schubert, P., Wilke, T., 2019. Impacts of microplastics on growth and health of hermatypic corals are species-specific. Environ. Pollut. 254, 113074. doi:10.1016/j.envpol.2019.113074

Reichert, J., Backes, A.R., Schubert, P., Wilke, T., 2017. The power of 3D fractal dimensions for comparative shape and structural complexity analyses of irregularly shaped organisms. Methods Ecol. Evol. 8, 1–9. doi:10.1111/2041-210X.12829

Reichert, J., Schellenberg, J., Schubert, P., Wilke, T., 2018. Responses of reef building corals to microplastic exposure. Environ. Pollut. 237, 955–960. doi:10.1016/j.envpol.2017.11.006

Reichert, J., Schellenberg, J., Schubert, P., Wilke, T., 2016. 3D scanning as a highly precise, reproducible, and minimally invasive method for surface area and volume measurements of scleractinian corals. Limnol. Oceanogr. Methods 14, 518–526. doi:10.1002/lom3.10109

Ripatti, S., Palmgren, J., 2000. Estimation of multivariate frailty models using penalized partial likelihood. Biometrics 56, 1016–1022. doi:10.1111/j.0006-341X.2000.01016.x

Rosic, N.N., Pernice, M., Dove, S., Dunn, S., Hoegh-Guldberg, O., 2011. Gene expression profiles of cytosolic heat shock proteins Hsp70 and Hsp90 from symbiotic dinoflagellates in response to thermal stress: Possible implications for coral bleaching. Cell Stress Chaperones 16, 69–80. doi:10.1007/s12192-010-0222-x

Rotjan, R.D., Sharp, K.H., Gauthier, A.E., Yelton, R., Lopez, E.M.B., Carilli, J., Kagan, J.C., Urban-Rich, J., 2019. Patterns, dynamics and consequences of microplastic ingestion by the temperate coral, Astrangia poculata. Proc. R. Soc. B Biol. Sci. 286, 20190726. doi:10.1098/rspb.2019.0726

Saliu, F., Montano, S., Leoni, B., Lasagni, M., Galli, P., 2019. Microplastics as a threat to coral reef environments: Detection of phthalate esters in neuston and scleractinian corals from the Faafu Atoll, Maldives. Mar. Pollut. Bull. 142, 234–241. doi:10.1016/j.marpolbul.2019.03.043

Savinelli, B., Vega Fernández, T., Galasso, N.M., D’Anna, G., Pipitone, C., Prada, F., Zenone, A., Badalamenti, F., Musco, L., 2020. Microplastics impair the feeding performance of a Mediterranean habitat-forming coral. Mar. Environ. Res. 155. doi:10.1016/j.marenvres.2020.104887

Seveso, D., Montano, S., Strona, G., Orlandi, I., Galli, P., Vai, M., 2014. The susceptibility of corals to thermal stress by analyzing Hsp60 expression. Mar. Environ. Res. 99, 69–75. doi:10.1016/j.marenvres.2014.06.008

Stafford, R., Jones, P.J.S., 2019. Viewpoint – Ocean plastic pollution: A convenient but distracting truth? Mar. Policy 103, 187–191. doi:10.1016/j.marpol.2019.02.003

Su, Y., Zhang, K., Zhou, Z., Wang, J., Yang, X., Tang, J., Li, H., Lin, S., 2020. Microplastic exposure represses the growth of endosymbiotic dinoflagellate Cladocopium goreaui in culture through affecting its apoptosis and metabolism. Chemosphere 244, 125485. doi:10.1016/j.chemosphere.2019.125485

Syakti, A.D., Jaya, J.V., Rahman, A., Hidayati, N.V., Raza’i, T.S., Idris, F., Trenggrono, M., Doumenq, P., Chou, L.M., 2019. Bleaching and necrosis of staghorn coral (Acropora formosa) in laboratory assays: Immediate impact of LDPE microplastics. Chemosphere 228, 528–535. doi:10.1016/j.chemosphere.2019.04.156

Tang, J., Ni, X., Zhou, Z., Wang, L., Lin, S., 2018. Acute microplastic exposure raises stress response and suppresses detoxification and immune capacities in the scleractinian coral Pocillopora damicornis. Environ. Pollut. 243, 66–74. doi:10.1016/j.envpol.2018.08.045

Tang, J., Wu, Z., Wan, L., Cai, W., Chen, S., Wang, X., Luo, J., Zhou, Z., Zhao, J., Lin, S., 2021. Differential enrichment and physiological impacts of ingested microplastics in scleractinian corals in situ. J. Hazard. Mater. 404, 124205. doi:10.1016/j.jhazmat.2020.124205

Tetu, S.G., Sarker, I., Schrameyer, V., Pickford, R., Elbourne, L.D.H., Moore, L.R., Paulsen, I.T., 2019. Plastic leachates impair growth and oxygen production in Prochlorococcus, the ocean’s most abundant photosynthetic bacteria. Commun. Biol. 2, 1–9. doi:10.1038/s42003-019-0410-x

Therneau, T.M., 2020. coxme: Mixed Effects Cox Models. R Packag. v 2.2-16.

Therneau, T.M., 1999. A package for survival analysis in S. R Packag. v 2.41-3 83.

Therneau, T.M., Grambsch, P.M., Pankratz, V.S., 2003. Penalized survival models and frailty. J. Comput. Graph. Stat. 12, 156–175. doi:10.1198/1061860031365

Veron, J.E.N., 2000. Corals of the World, 3rd ed. Australia: Australian Institute of Marine Sciences and CRR Qld Pty Ltd.

Voolstra, C.R., Buitrago-López, C., Perna, G., Cárdenas, A., Hume, B.C.C., Rädecker, N., Barshis, D.J., 2020. Standardized short-term acute heat stress assays resolve historical differences in coral thermotolerance across microhabitat reef sites. Glob. Chang. Biol. 26, 4328–4343. doi:10.1111/gcb.15148

Wickham, H., 2016. ggplot2: Elegant Graphics for Data Analysis. Springer-Verlag New York.

Wiedenmann, J., D’Angelo, C., Smith, E.G., Hunt, A.N., Legiret, F.E., Postle, A.D., Achterberg, E.P., 2013. Nutrient enrichment can increase the susceptibility of reef corals to bleaching. Nat. Clim. Chang. 3, 160–164. doi:10.1038/nclimate1661

