## Supplementary information for "Interactive effects of microplastic pollution and heat stress on reef-building corals"

### Supplementary figures

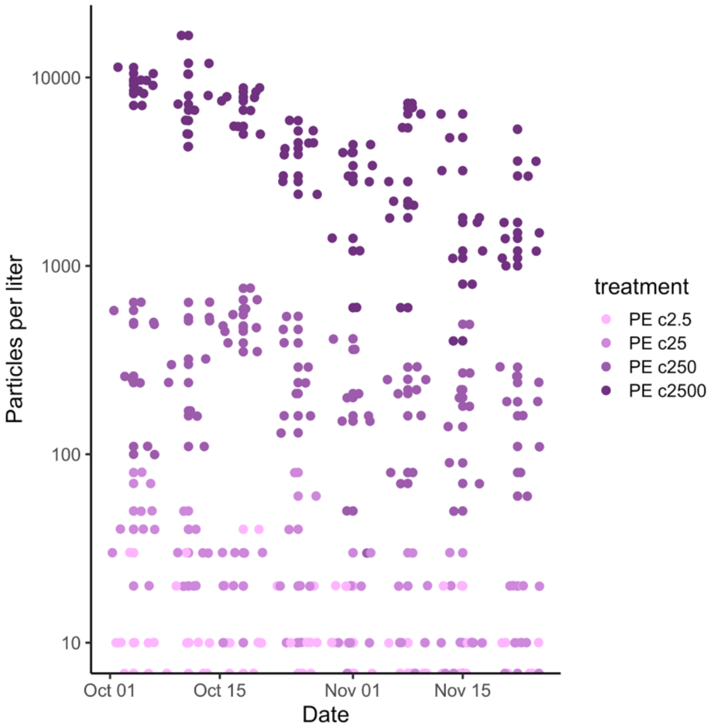


**Supplementary Figure 1.** Concentration of PE microplastic particles in replicate tanks over nine weeks experimental exposure of the stony coral species *Pocillopora verrucosa* and *Stylophora pistillata* (Experiment 1).

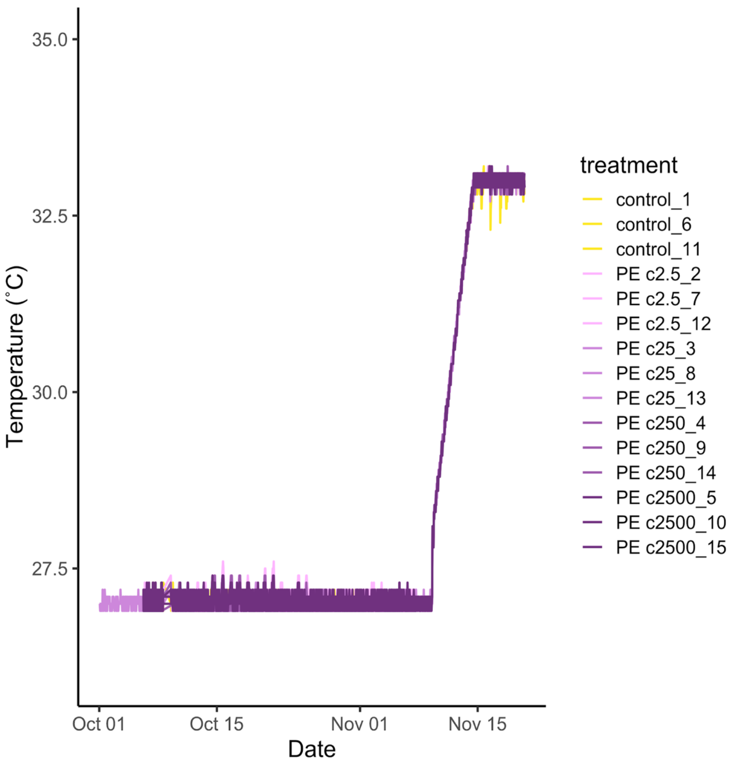


**Supplementary Figure 2.** Water temperature profile in replicate treatments of different concentrations of PE microplastic particles over nine weeks exposure of the stony coral species *Pocillopora verrucosa* and *Stylophora pistillata* (Experiment 1).

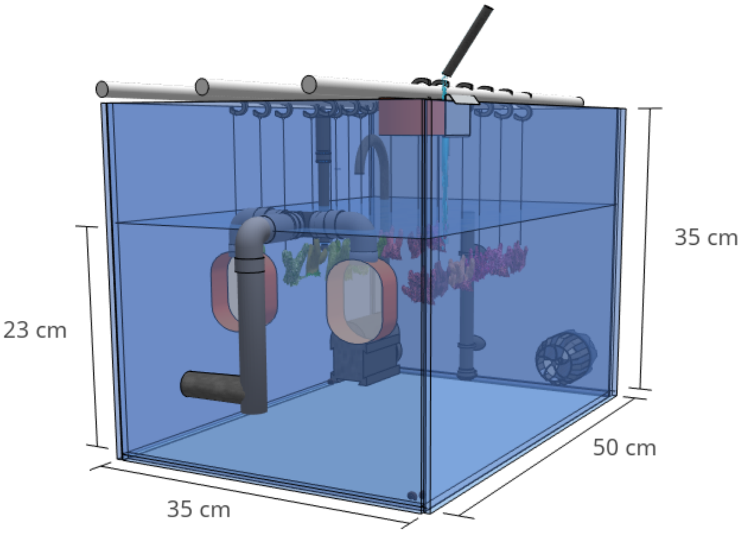


**Supplementary Figure 3.** 3D-model of an experimental tank with hanging coral fragments of the stony coral species *Pocillopora verrucosa* and *Stylophora pistillata* (Experiment 1 & 3); the different components of the set up include: water inflow, inflow filter, temperature sensor, heater, turnover pump, current pump, outflow filter, outflow.

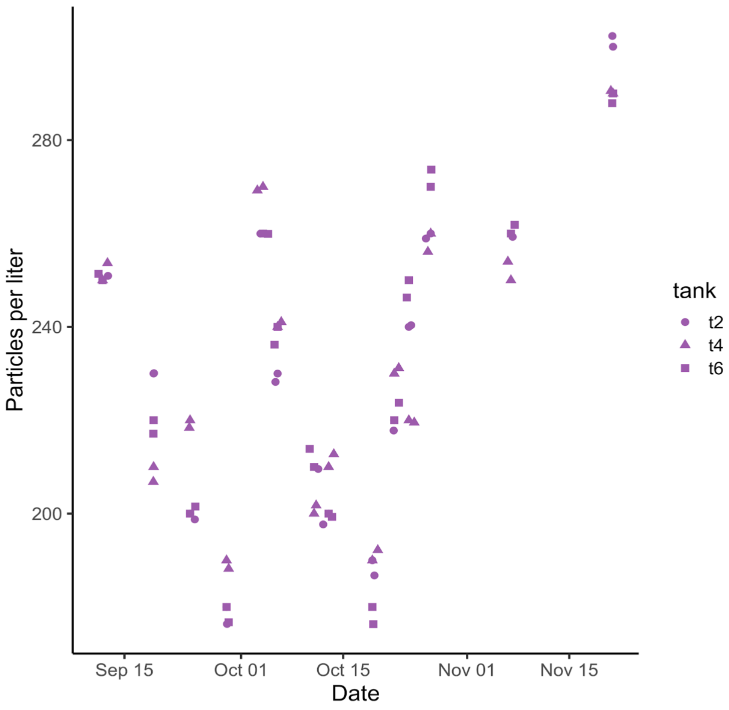


**Supplementary Figure 4.** Concentration of PE microplastic particles in replicate tanks over 15 weeks experimental exposure of the five stony coral species *Acropora muricata*, *Montipora digitata*, *Porites cylindrica*, *Pocillopora verrucosa*, and *Stylophora pistillata* (Experiment 2).

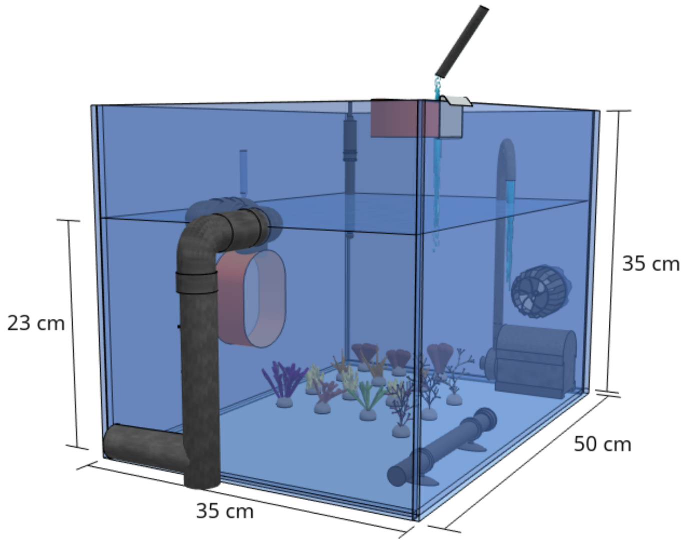


**Supplementary Figure 5.** 3D-model of an experimental tank with coral fragments of the species *Acropora muricata*, *Montipora digitata*, *Porites cylindrica*, *Pocillopora verrucosa*, and *Stylophora pistillata* in the center (Experiment 2); the components of the set up include: temperature sensor, inflow filter (mesh size 150 µm), water inflow, current pump, turnover pump, outflow filter (mesh size 65 µm), water outflow, and heater.

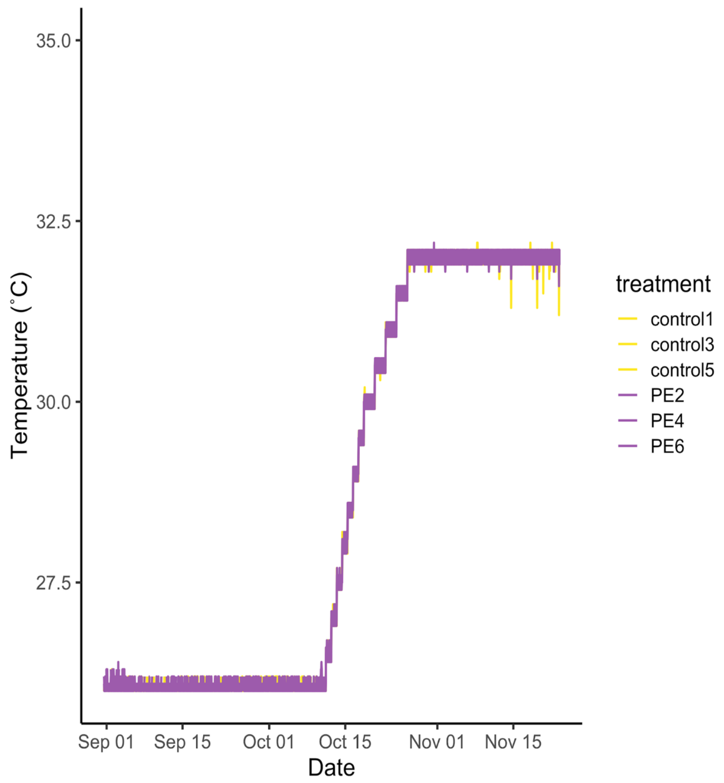


**Supplementary Figure 6.** Water temperature profile in replicate tanks over 15 weeks experimental exposure of the five stony coral species *Acropora muricata*, *Montipora digitata*, *Porites cylindrica*, *Pocillopora verrucosa*, and *Stylophora pistillata* (Experiment 2).

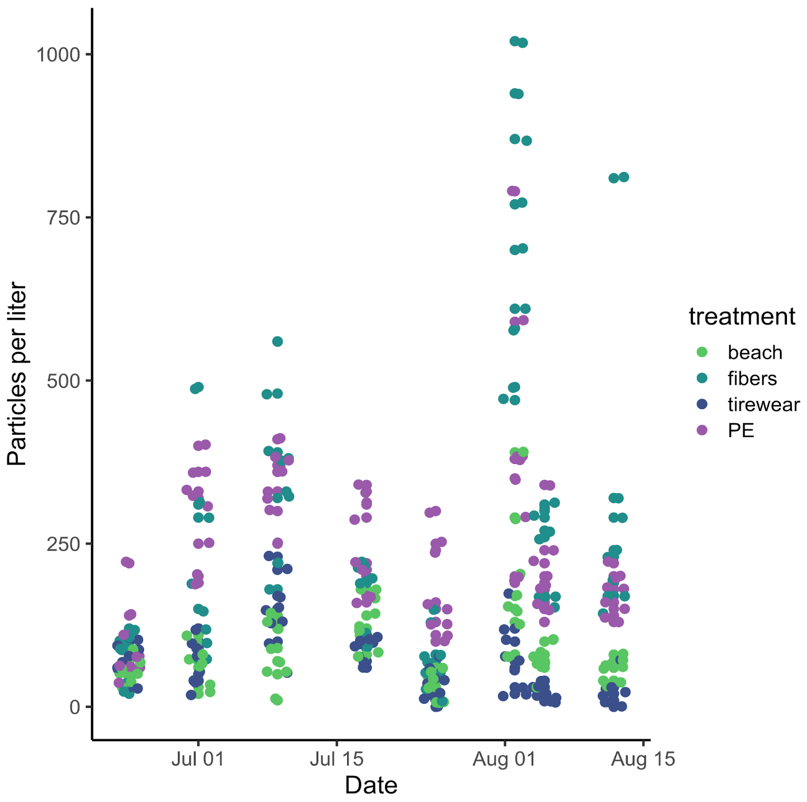


**Supplementary Figure 7.** Concentration of different types of microplastic particles over 10 weeks experimental exposure of the stony coral species *Pocillopora verrucosa* and *Stylophora pistillata* (Experiment 3).

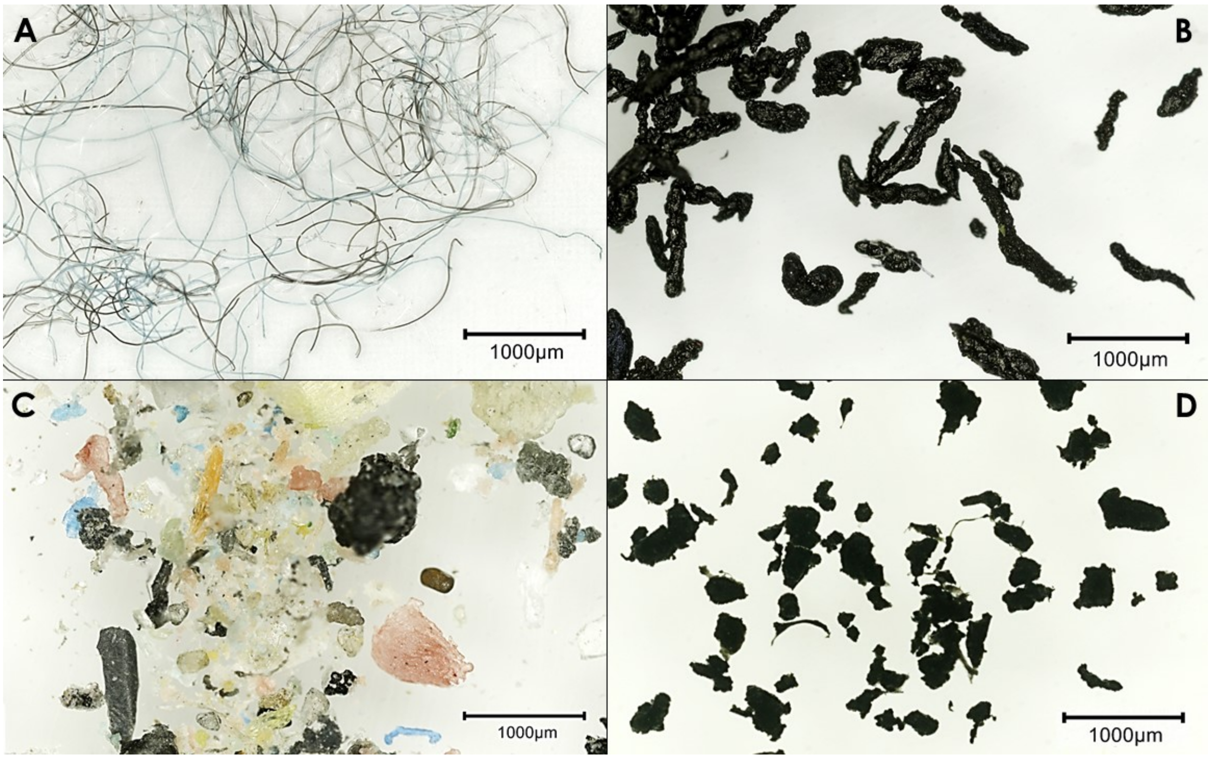


**Supplementary Figure 8.** Microparticles used for treatments in Experiment 3. A) artificial fibers from the fashion industry (treatment: “fibers”), B) combination of residues from the automobile sector consisting of tire wear, brake abrasion, and varnish (treatment: “tirewear”), C) mixture of secondary marine microplastics from fragmented plastic debris (treatment: “beach”), D) high-density polyethylene particles (treatment: “PE”).

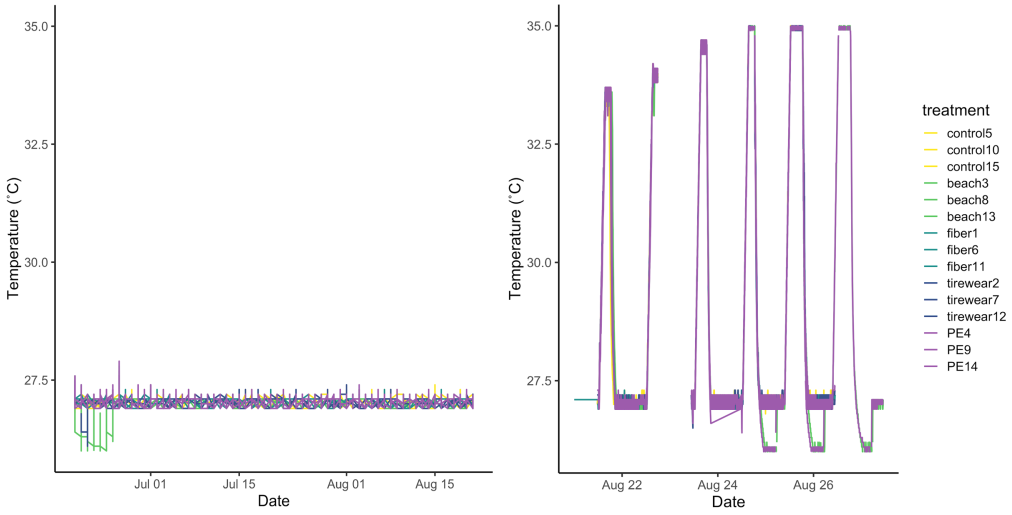


**Supplementary Figure 9.** Water temperature profile in different types of microplastic treatments over 10 weeks experimental exposure of the stony coral species *Pocillopora verrucosa* and *Stylophora pistillata* (Experiment 3). The experiment was divided into a stable phase (left) and a one-week period at the end of repeated heat stress treatments (right).

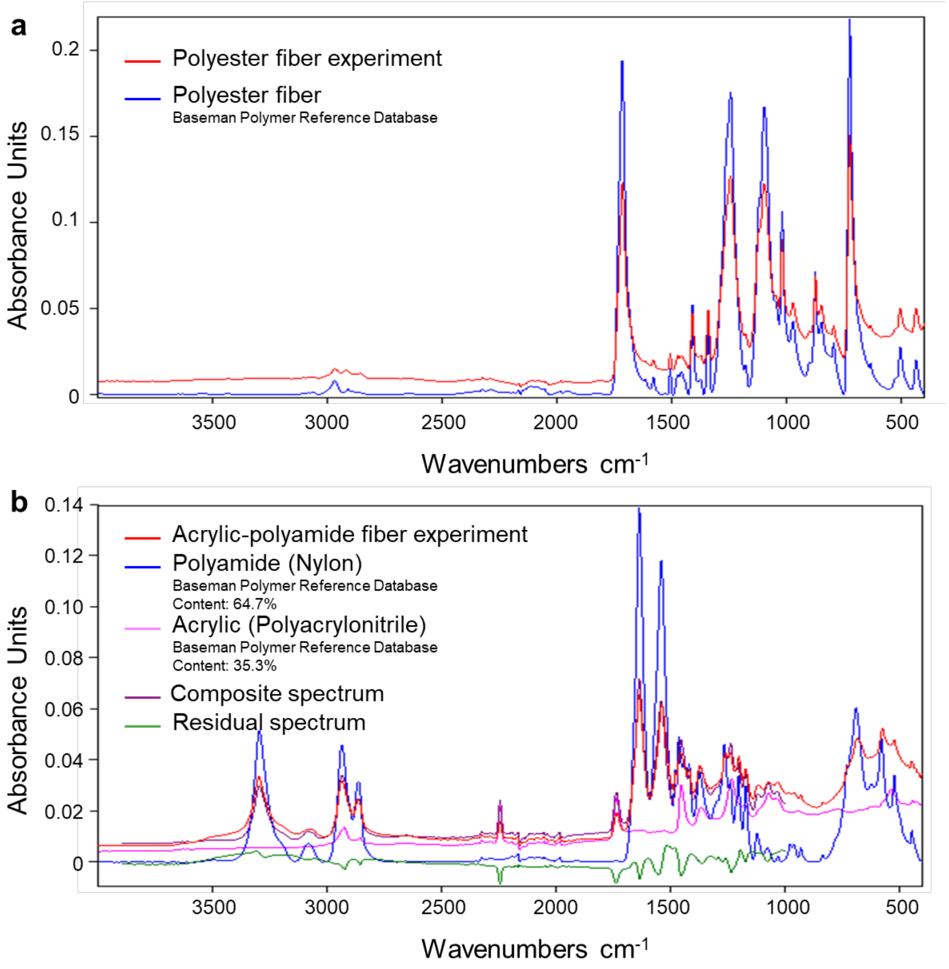


**Supplementary Figure 10.** FTIR spectrum of polyester (a) and acrylic-polyamide fibers used in the experiment (red), derived from ATR-FTIR and compared to reference spectra from the Baseman Polymer reference Database (Primpke et al., 2018).

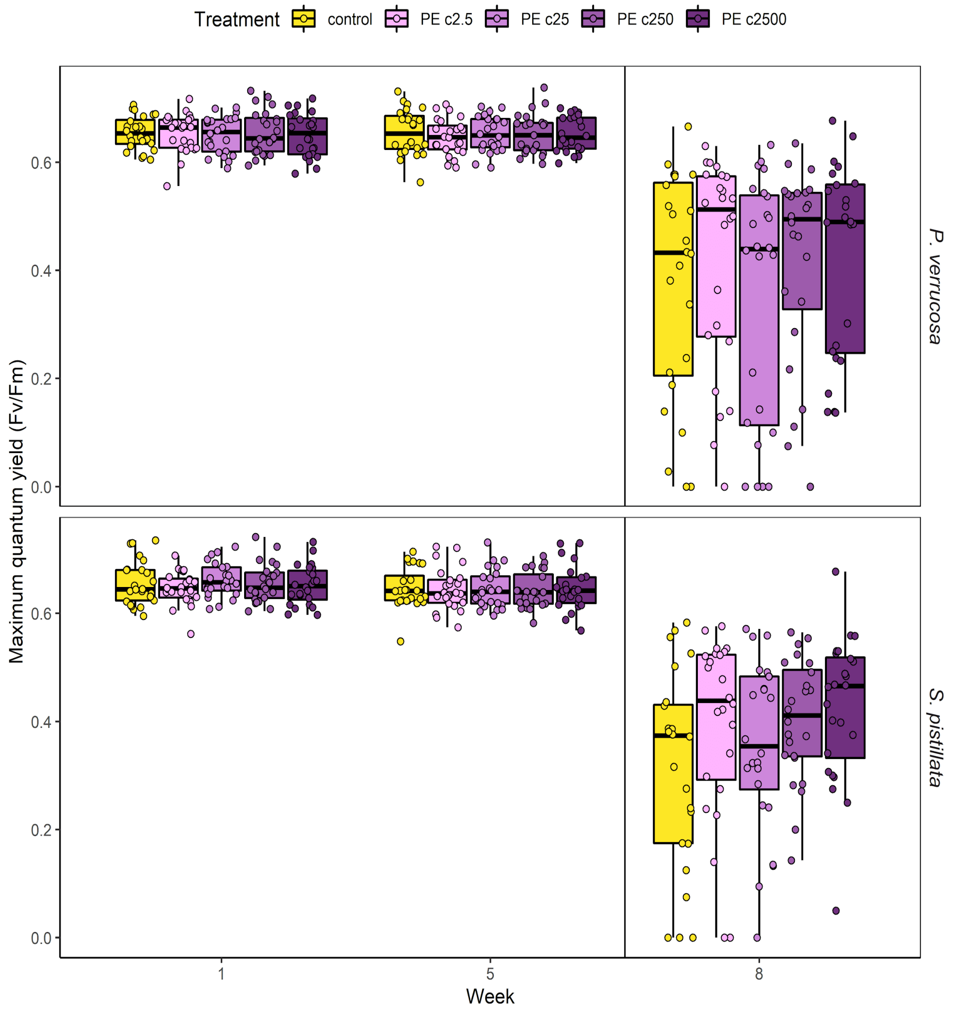


**Supplementary Figure 11.** Maximum photosynthetic efficiency (Fv/Fm) of the stony coral species *Pocillopora verrucosa* and *Stylophora pistillata* treated with different concentrations of PE microplastic particles and a microplastic-free control. The vertical line marks the onset of an additional heat treatment to test the effect of microplastic exposure on coral heat tolerance. Different concentrations of PE microplastic particles were compared to a microplastic-free control (heat only). Treatments of c2.5, c25, c250, c2500 indicate 2.5, 25, 250, 2 500 particles $L^{-1}$, respectively. Fv/Fm measured using pulse-amplitude-modulated fluorometry, with decreasing values indicating lower photosynthetic efficiency (photodamage) of the coral-associated photosymbionts.

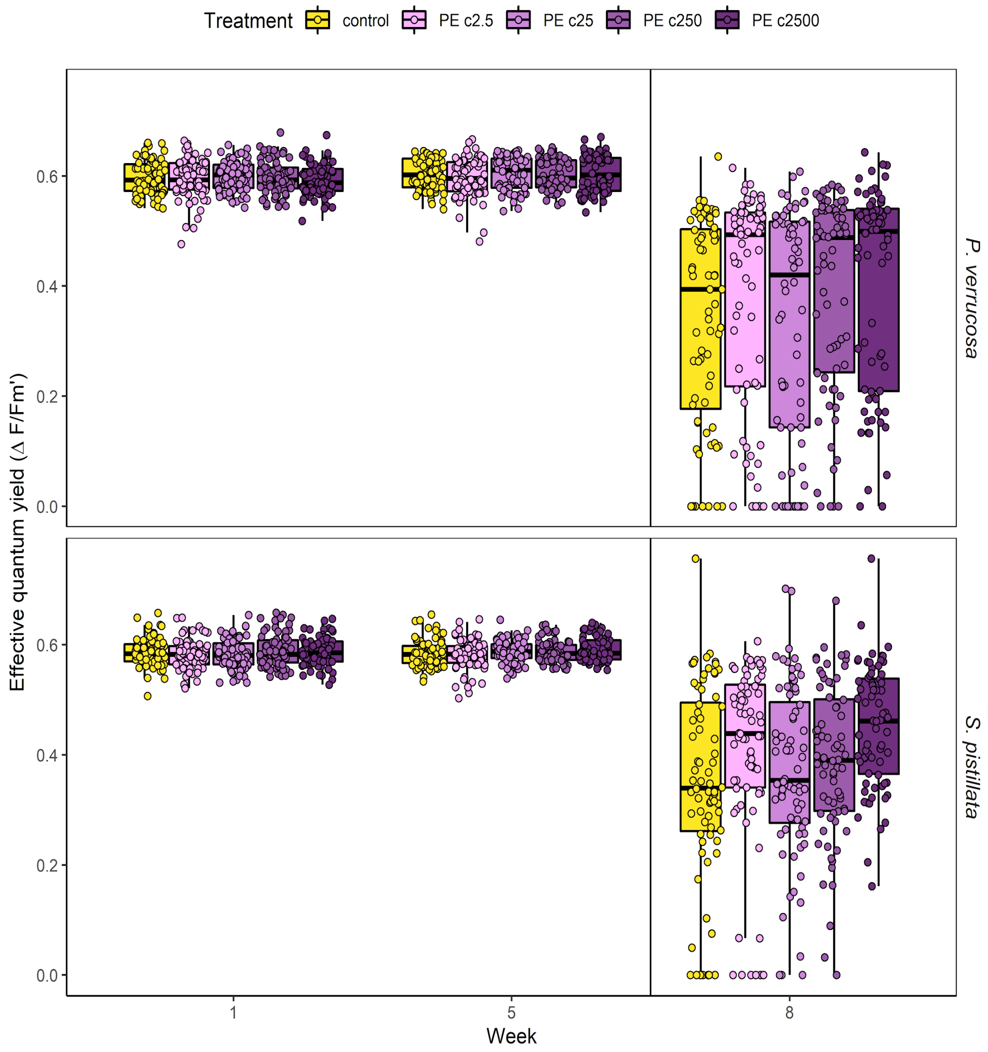


**Supplementary Figure 12.** Effective photosynthetic efficiency (ΔF/Fm') of the stony coral species *Pocillopora verrucosa* and *Stylophora pistillata* treated with different concentrations of PE microplastic particles and a microplastic-free control. The vertical line marks the onset of an additional heat treatment to test the effect of microplastic exposure on coral heat tolerance. Different concentrations of PE microplastic particles were compared to a microplastic-free control (heat only). Treatments of c2.5, c25, c250, c2500 indicate 2.5, 25, 250, 2 500 particles $L^{-1}$, respectively. ΔF/Fm' measured using pulse-amplitude-modulated fluorometry, with decreasing values indicating lower photosynthetic efficiency (photoinhibition) of the coral-associated photosymbionts.

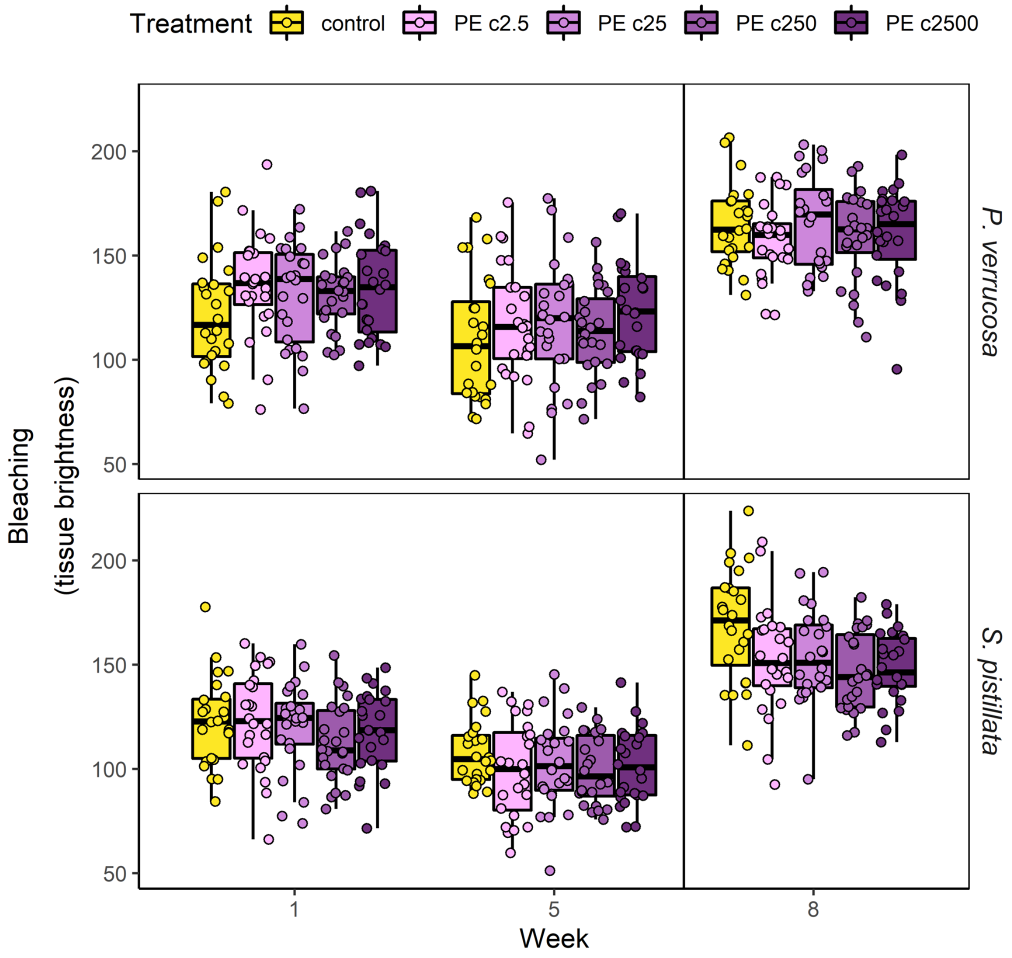


**Supplementary Figure 13.** Coral bleaching expressed as tissue brightness of the stony coral species *Pocillopora verrucosa* and *Stylophora pistillata* treated with different concentrations of PE microplastic particles and a microplastic-free control. The vertical line marks the onset of an additional heat treatment to test the effect of microplastic exposure on coral heat tolerance. Different concentrations of PE microplastic particles were compared to a microplastic-free control (heat only). Treatments of c2.5, c25, c250, c2500 indicate 2.5, 25, 250, 2 500 particles $L^{-1}$, respectively. Brightness values derived from standardized photographs. Brightness expressed as pixel values: 0 = black, 255 = white, i.e. the larger the number, the brighter the coral fragment.

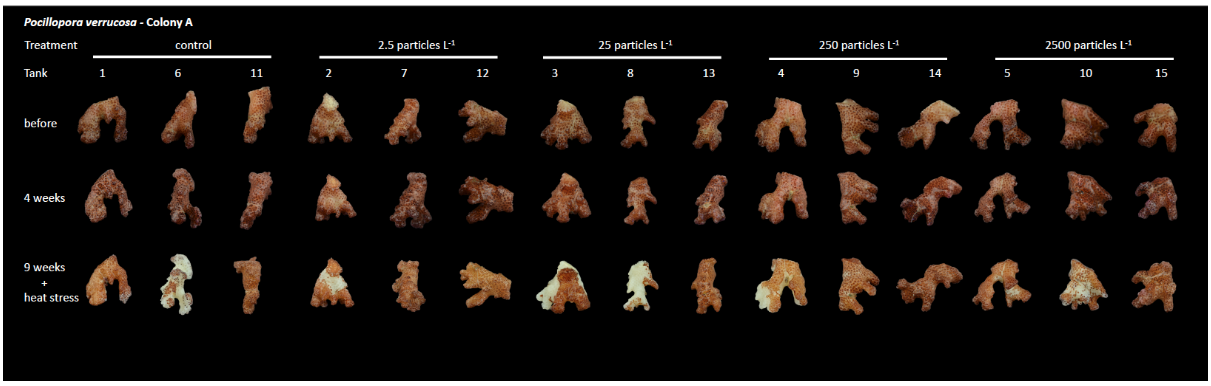


**Supplementary Figure 14.** Overview of all fragments of colony A of the stony coral species *Pocillopora verrucosa* before the experimental treatment, at 4 weeks of microplastic exposure, and at 9 weeks of microplastic exposure in combination with heat stress. Pictures were used to calculate brightness as a measure of coral bleaching.

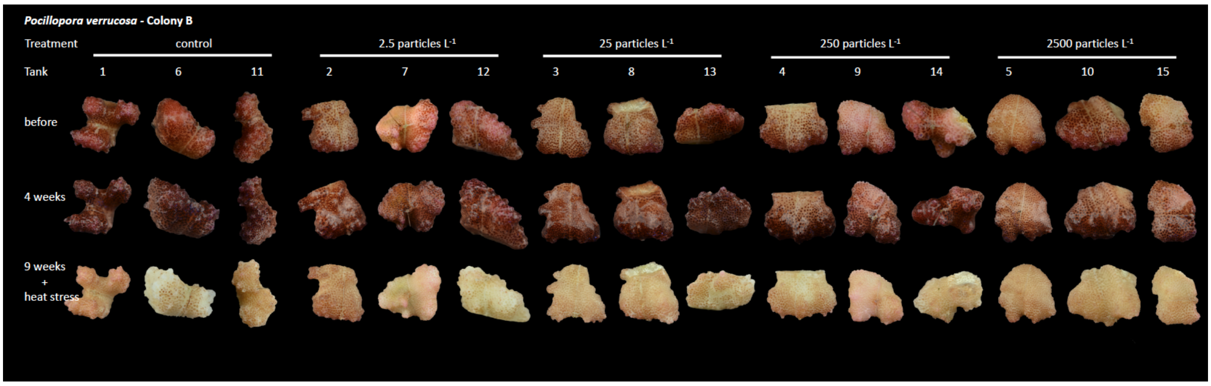


**Supplementary Figure 15.** Overview of all fragments of colony B of the stony coral species *Pocillopora verrucosa* before the experimental treatment, at 4 weeks of microplastic exposure, and at 9 weeks of microplastic exposure in combination with heat stress. Pictures were used to calculate brightness as a measure of coral bleaching.

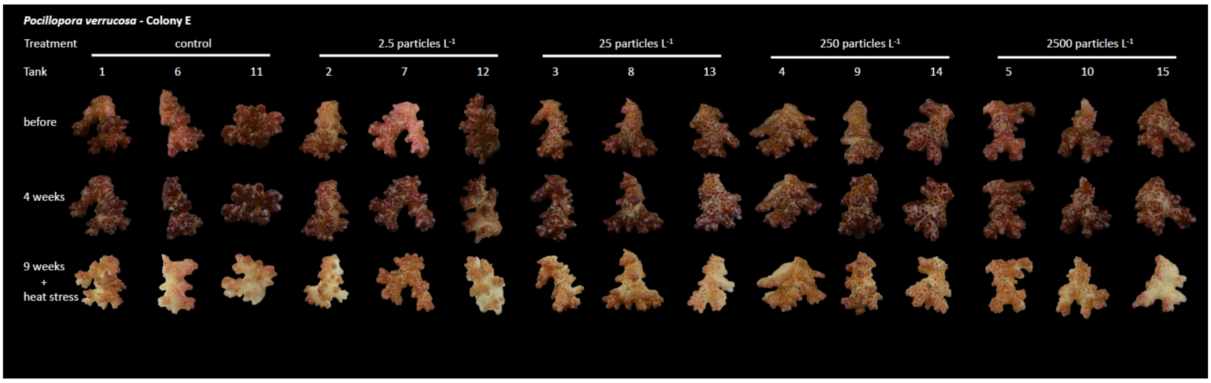


**Supplementary Figure 16.** Overview of all fragments of colony E of the stony coral species *Pocillopora verrucosa* before the experimental treatment, at 4 weeks of microplastic exposure, and at 9 weeks of microplastic exposure in combination with heat stress. Pictures were used to calculate brightness as a measure of coral bleaching.

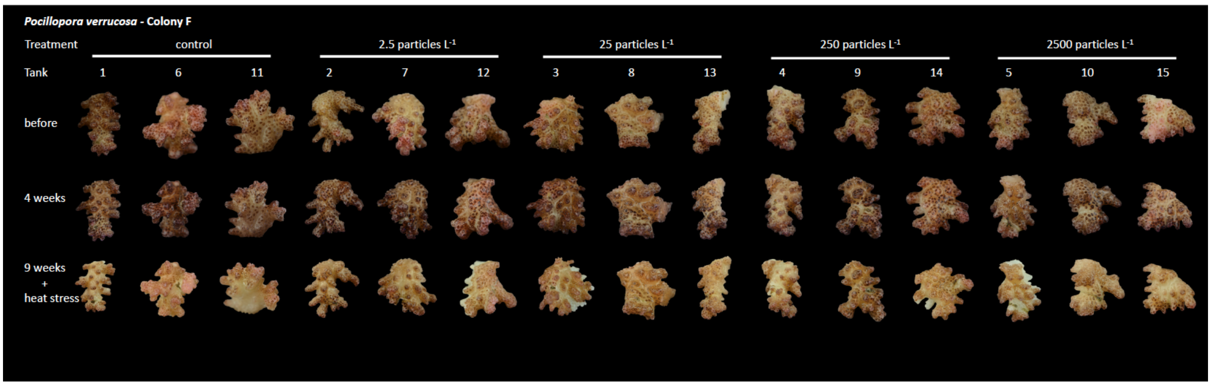


**Supplementary Figure 17.** Overview of all fragments of colony F of the stony coral species *Pocillopora verrucosa* before the experimental treatment, at 4 weeks of microplastic exposure, and at 9 weeks of microplastic exposure in combination with heat stress. Pictures were used to calculate brightness as a measure of coral bleaching.

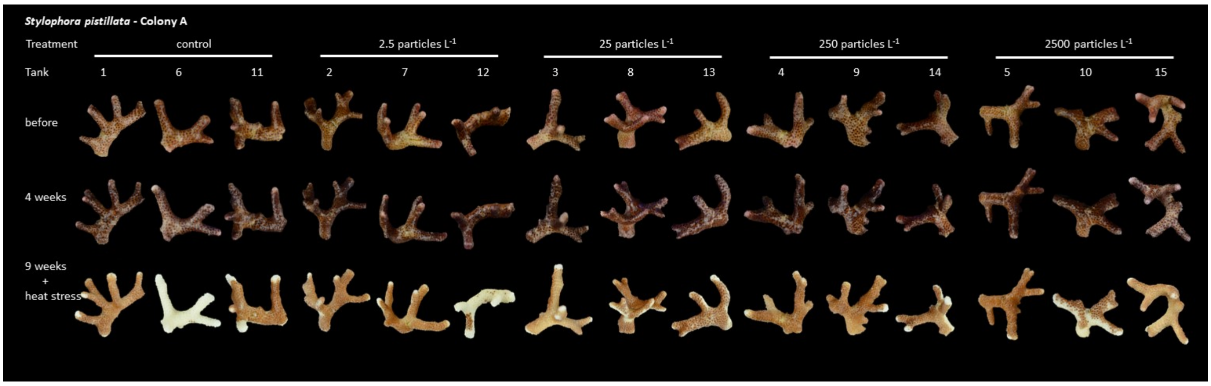


**Supplementary Figure 18.** Overview of all fragments of colony A of the stony coral species *Stylophora pistillata* before the experimental treatment, at 4 weeks of microplastic exposure, and at 9 weeks of microplastic exposure in combination with heat stress. Pictures were used to calculate brightness as a measure of coral bleaching.

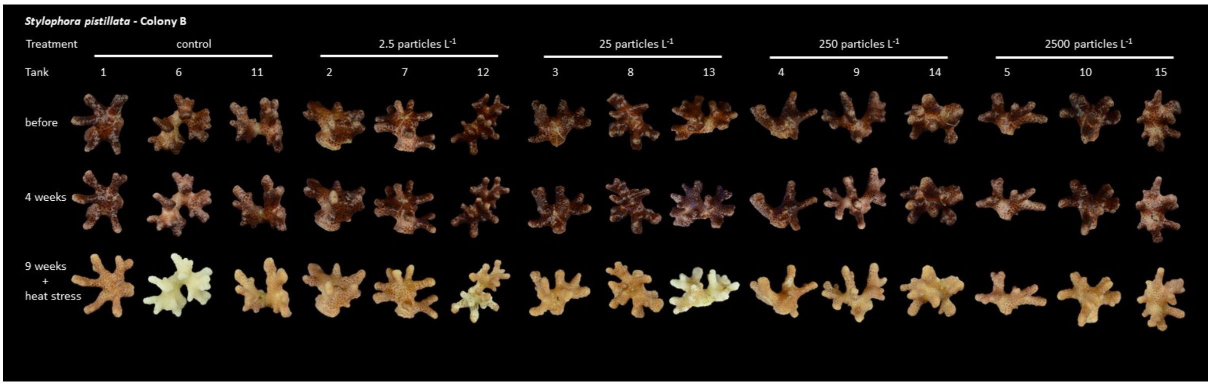


**Supplementary Figure 19.** Overview of all fragments of colony B of the stony coral species *Stylophora pistillata* before the experimental treatment, at 4 weeks of microplastic exposure, and at 9 weeks of microplastic exposure in combination with heat stress. Pictures were used to calculate brightness as a measure of coral bleaching.

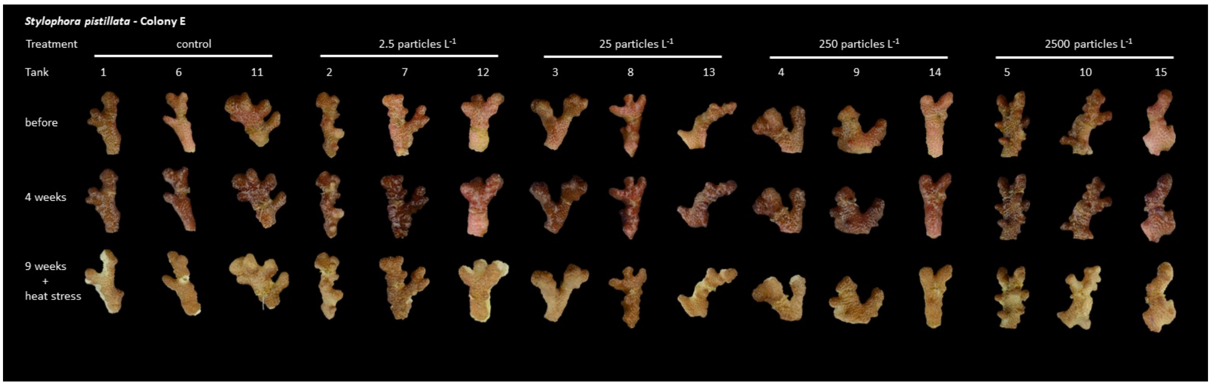


**Supplementary Figure 20.** Overview of all fragments of colony E of the stony coral species *Stylophora pistillata* before the experimental treatment, at 4 weeks of microplastic exposure, and at 9 weeks of microplastic exposure in combination with heat stress. Pictures were used to calculate brightness as a measure of coral bleaching.

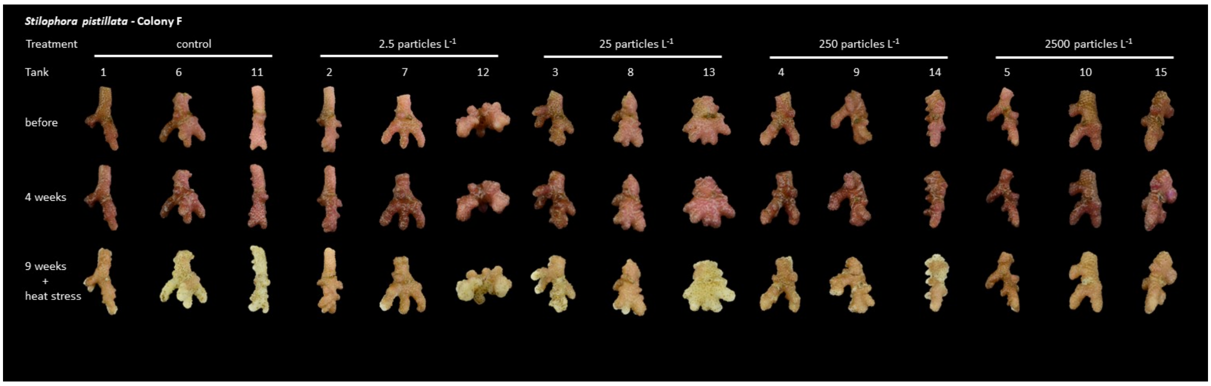


**Supplementary Figure 21.** Overview of all fragments of colony F of the stony coral species *Stylophora pistillata* before the experimental treatment, at 4 weeks of microplastic exposure, and at 9 weeks of microplastic exposure in combination with heat stress. Pictures were used to calculate brightness as a measure of coral bleaching.

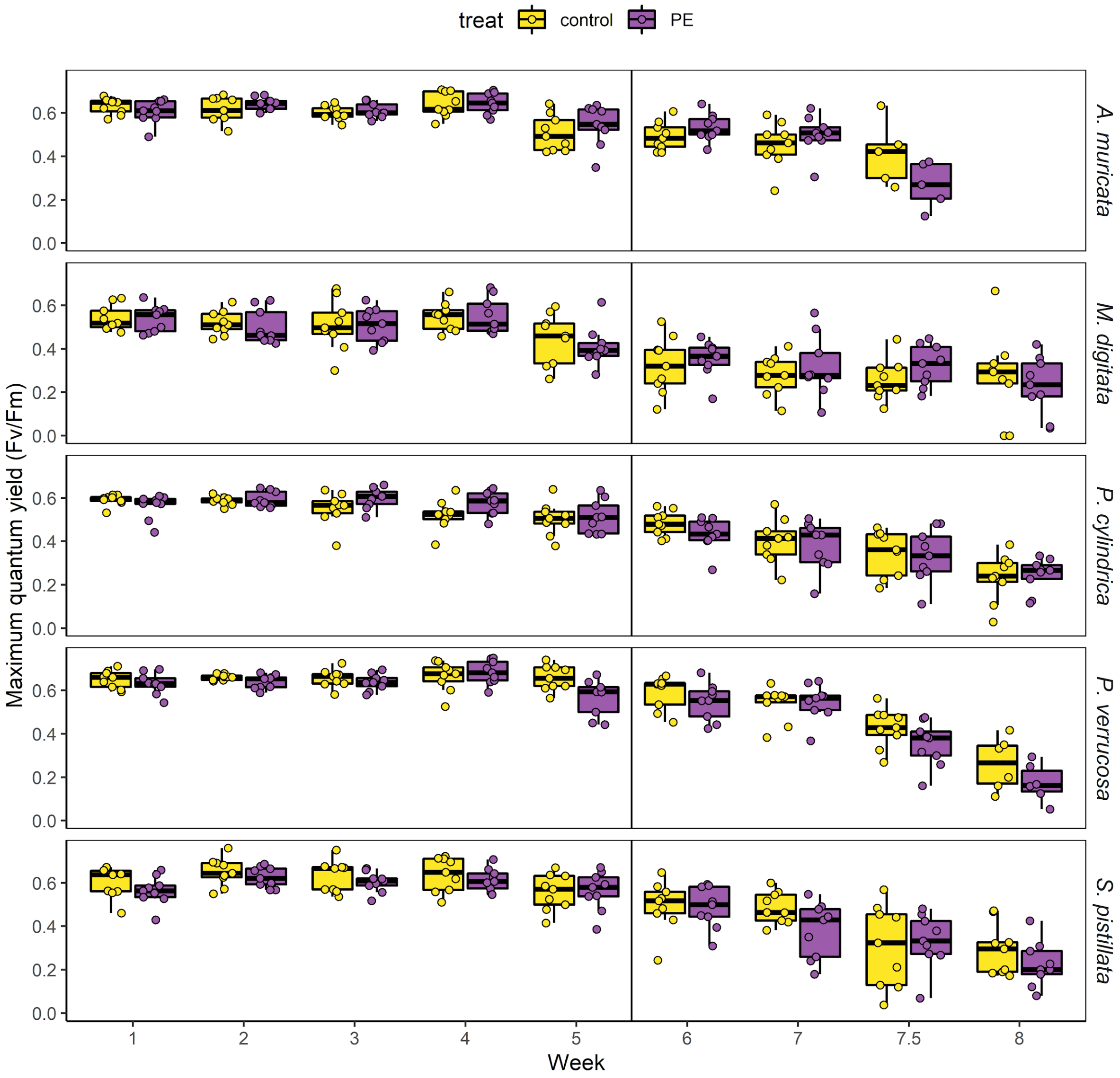


**Supplementary Figure 22.** Maximum photosynthetic efficiency (Fv/Fm) of stony coral fragments from the five species *Acropora muricata*, *Montipora digitata*, *Porites cylindrica*, *Pocillopora verrucosa*, and *Stylophora pistillata* treated with PE microplastic particles and a microplastic-free control. The vertical line marks the onset of an additional heat treatment to test the effect of microplastic exposure on coral heat tolerance. Fv/Fm measured using pulse-amplitude-modulated fluorometry, with decreasing values indicating lower photosynthetic efficiency (photodamage) of the coral-associated photosymbionts.

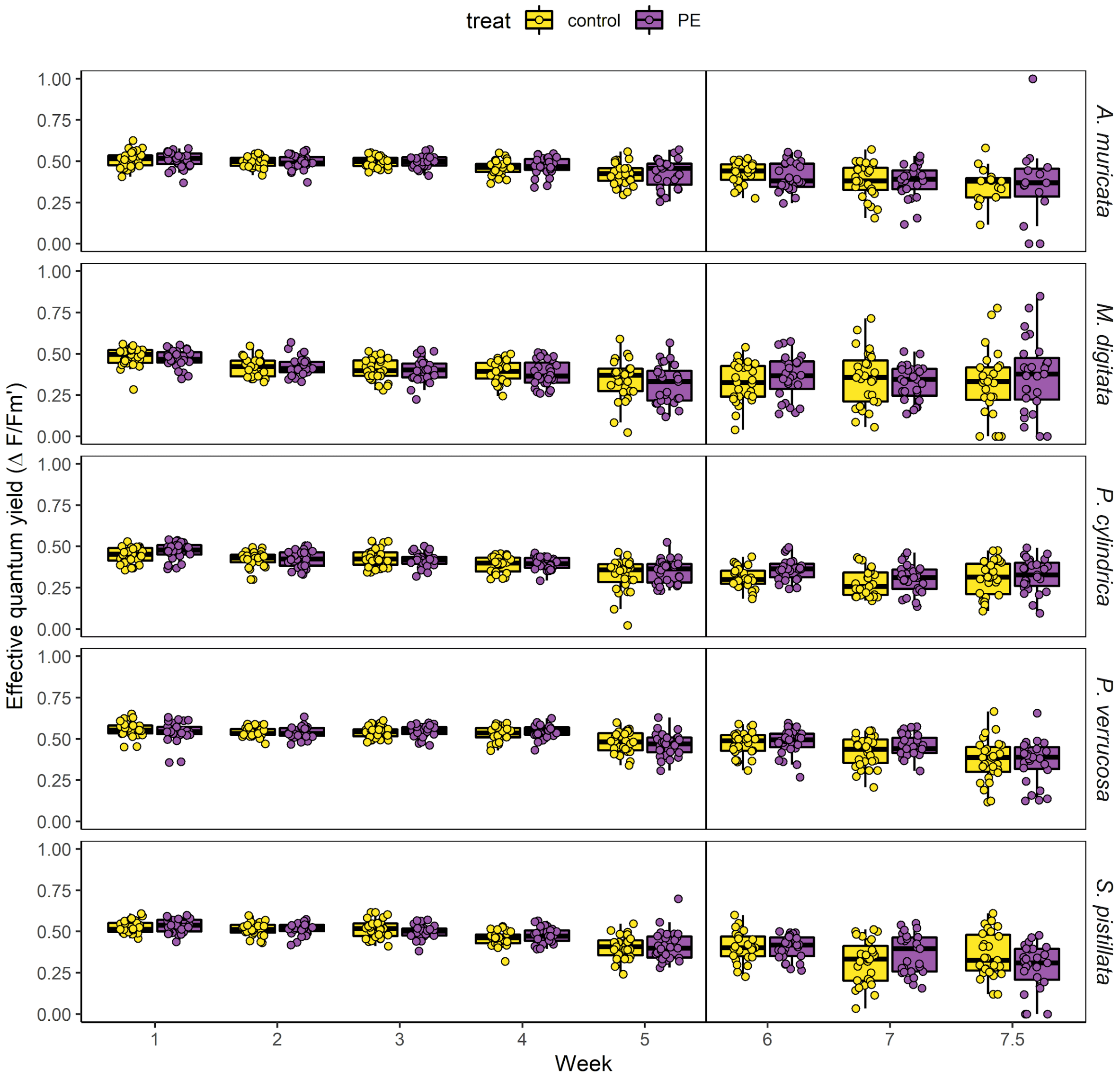


**Supplementary Figure 23.** Effective photosynthetic efficiency (ΔF/Fm') of stony coral fragments from five species *Acropora muricata*, *Montipora digitata*, *Porites cylindrica*, *Pocillopora verrucosa*, and *Stylophora pistillata* treated with PE microplastic particles and a microplastic-free control. The vertical line marks the onset of an additional heat treatment to test the effect of microplastic exposure on coral heat tolerance. ΔF/Fm' measured using pulse-amplitude-modulated fluorometry, with decreasing values indicating lower photosynthetic efficiency (photoinhibition) of the coral-associated photosymbionts.

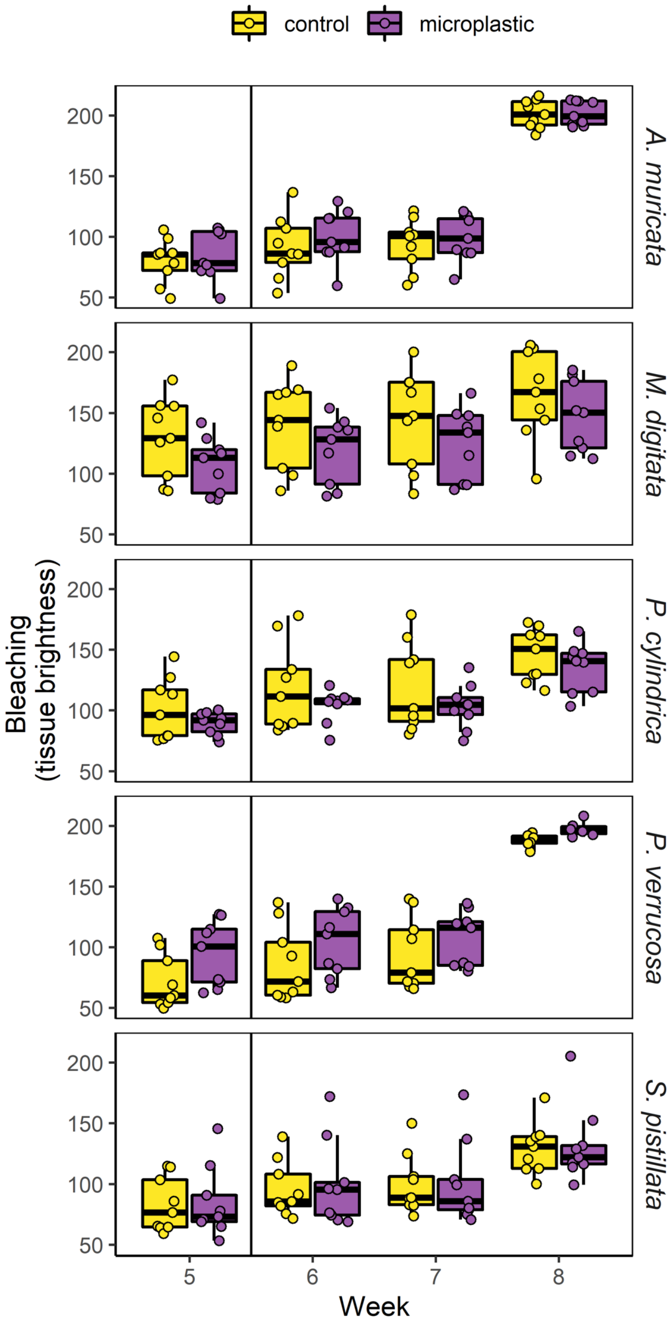


**Supplementary Figure 24.** Coral bleaching expressed as tissue brightness of stony coral fragments from the five *Acropora muricata*, *Montipora digitata*, *Porites cylindrica*, *Pocillopora verrucosa*, and *Stylophora pistillata* species treated with PE microplastic particles and a microplastic-free control. The vertical line marks the onset of an additional heat treatment to test the effect of microplastic exposure on coral heat tolerance. Brightness values derived from standardized photographs. Brightness expressed as pixel values: 0 = black, 255 = white, i.e. the larger the number, the brighter the coral fragment.

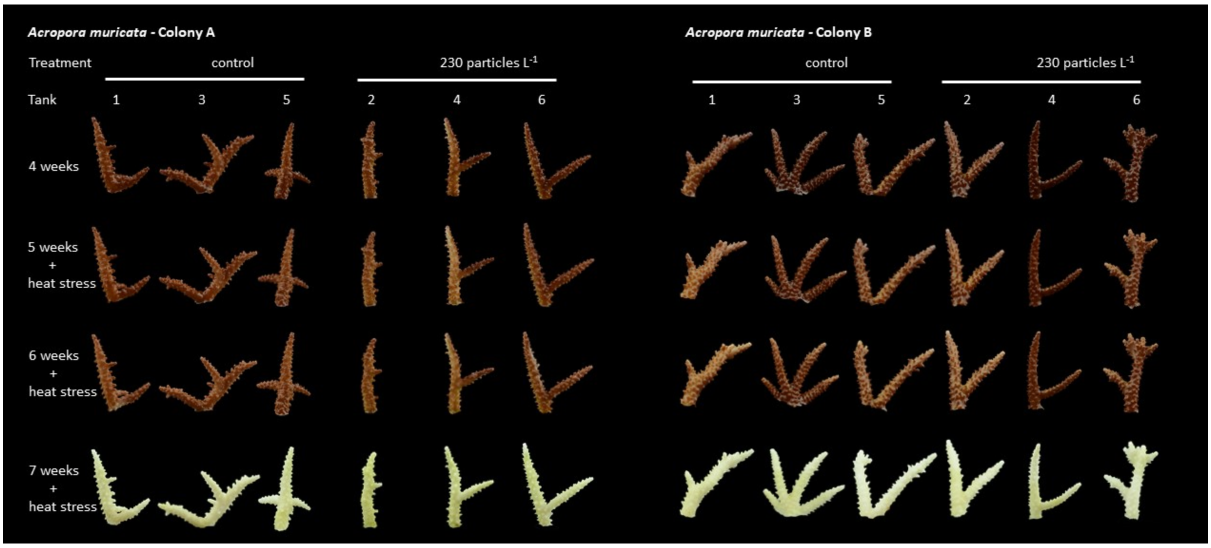


**Supplementary Figure 25.** Overview of all fragments of the stony coral species *Acropora muricata* after 4, 5, 6, and 7 weeks experimental treatment. Corals were exposed to microplastic or a microplastic-free control for 4 weeks, after which all experimental groups were exposed to additional heat stress. Pictures were used to calculate brightness as a measure of coral bleaching.

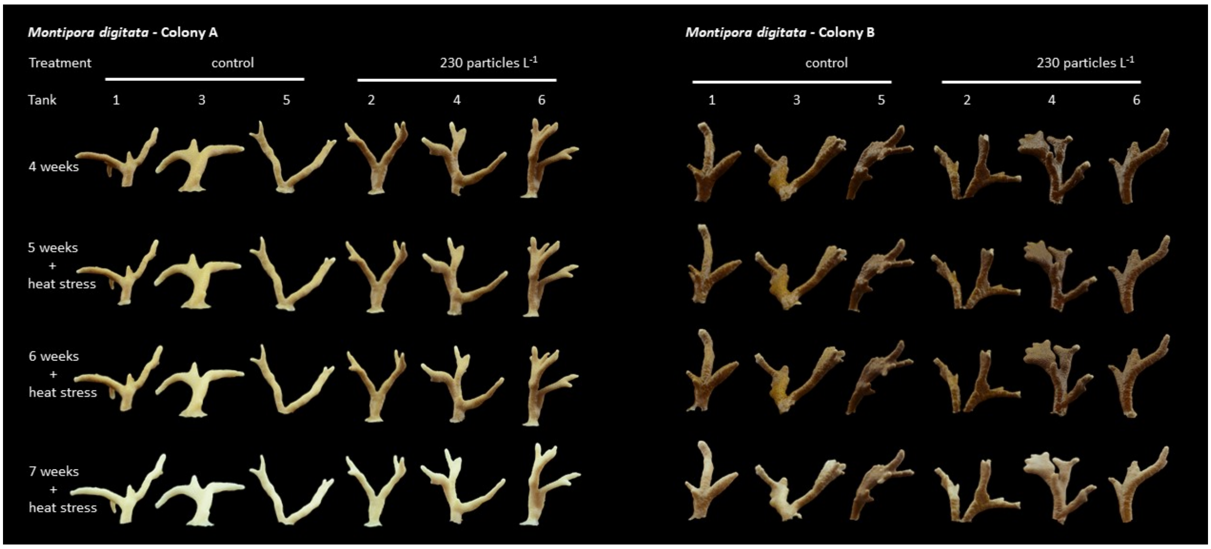


**Supplementary Figure 26.** Overview of all fragments of the stony coral species *Montipora digitata* after 4, 5, 6, and 7 weeks experimental treatment. Corals were exposed to microplastic or a microplastic-free control for 4 weeks, after which all experimental groups were exposed to additional heat stress. Pictures were used to calculate brightness as a measure of coral bleaching.

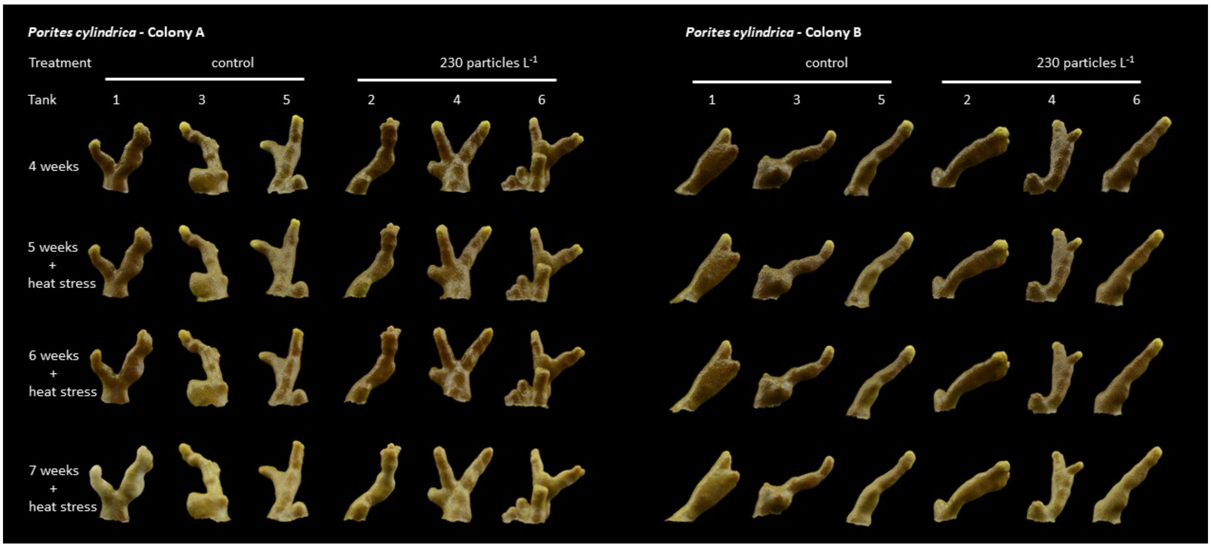


**Supplementary Figure 27.** Overview of all fragments of the stony coral species *Porites cylindrica* after 4, 5, 6, and 7 weeks experimental treatment. Corals were exposed to microplastic or a microplastic-free control for 4 weeks, after which all experimental groups were exposed to additional heat stress. Pictures were used to calculate brightness as a measure of coral bleaching.

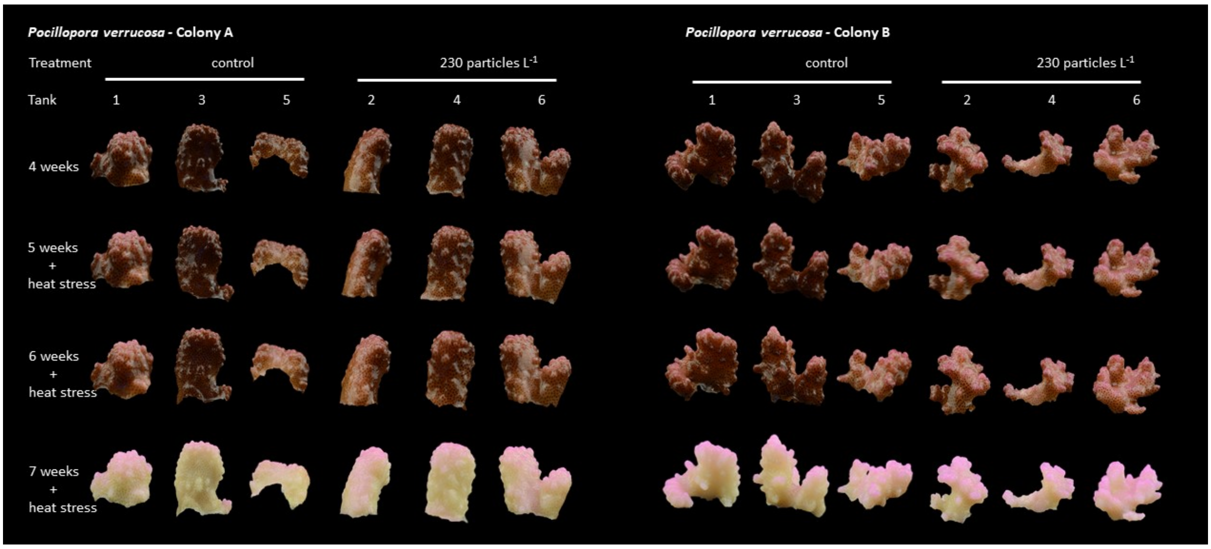


**Supplementary Figure 28.** Overview of all fragments of the stony coral species *Pocillopora verrucosa* after 4, 5, 6, and 7 weeks experimental treatment. Corals were exposed to microplastic or a microplastic-free control for 4 weeks, after which all experimental groups were exposed to additional heat stress. Pictures were used to calculate brightness as a measure of coral bleaching.

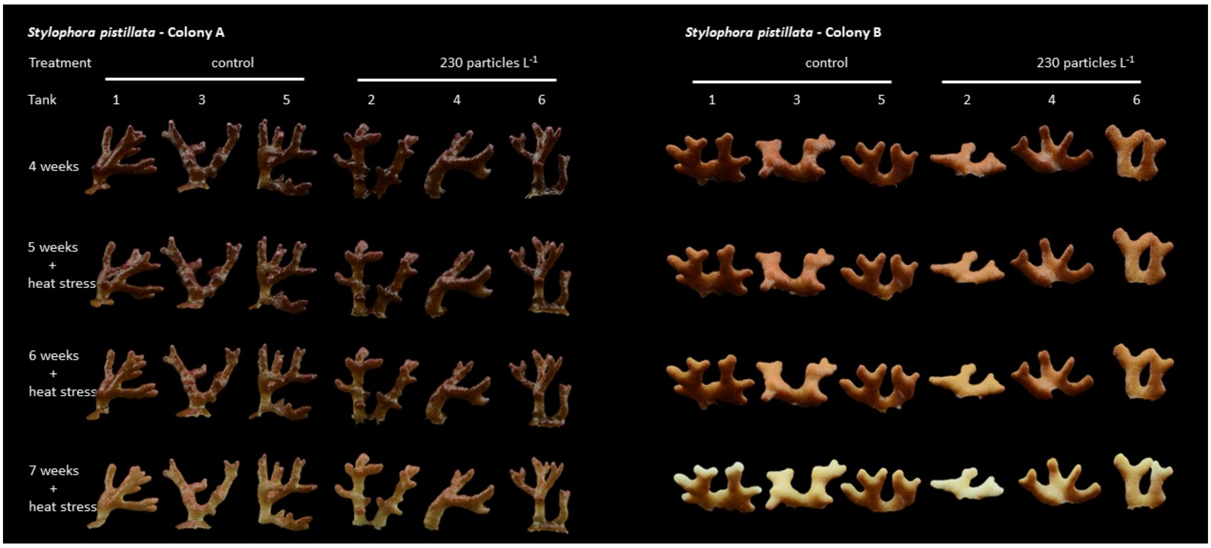


**Supplementary Figure 29.** Overview of all fragments of the stony coral species.*Stylophora pistillata* after 4, 5, 6, and 7 weeks experimental treatment. Corals were exposed to microplastic or a microplastic-free control for 4 weeks, after which all experimental groups were exposed to additional heat stress. Pictures were used to calculate brightness as a measure of coral bleaching.

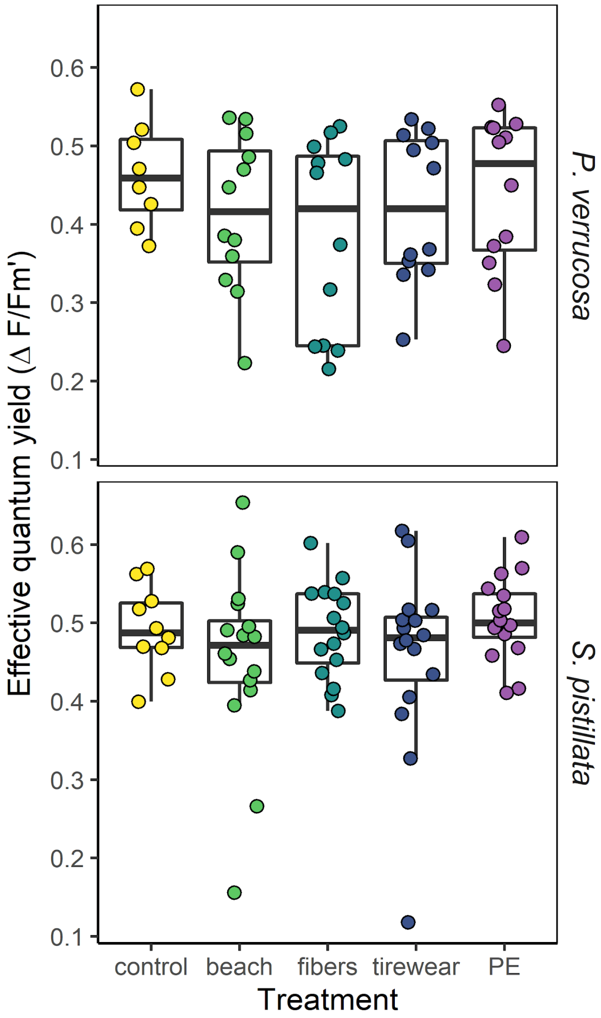


**Supplementary Figure 30.** Effective photosynthetic efficiency (ΔF/Fm') of fragments of the stony coral species *Pocillopora verrucosa* and *Stylophora pistillata* after 10 weeks of combined microplastic and heat stress treatment. Different types of microplastic particles were compared to a microplastic-free control (heat only). ΔF/Fm' measured using pulse-amplitude-modulated fluorometry, with decreasing values indicating lower photosynthetic efficiency (photoinhibition) of the coral-associated photosymbionts.

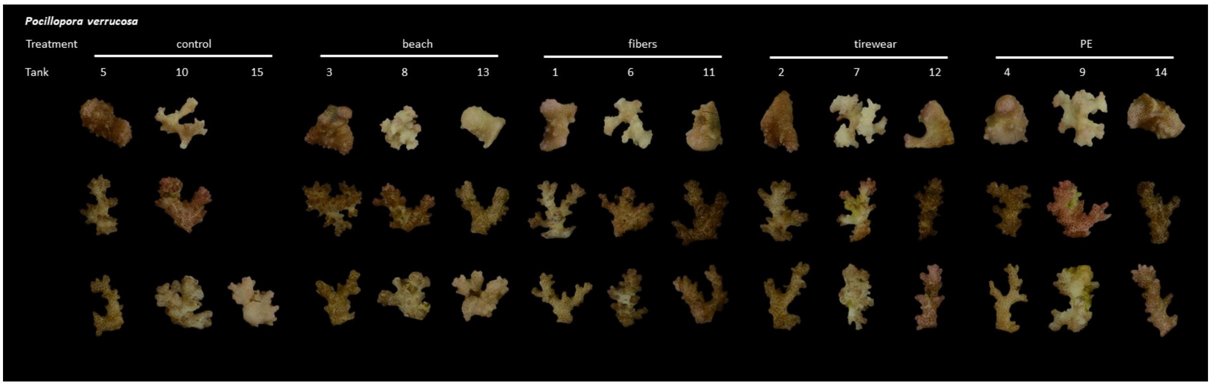


**Supplementary Figure 31.** Overview of all fragments of the stony coral species *Pocillopora verrucosa* after 10 weeks of combined microplastic and heat stress treatment. The displayed pictures were used to calculate brightness as a measure of coral bleaching. Fragments from tank 15 are missing due to a technical error in the aquarium.

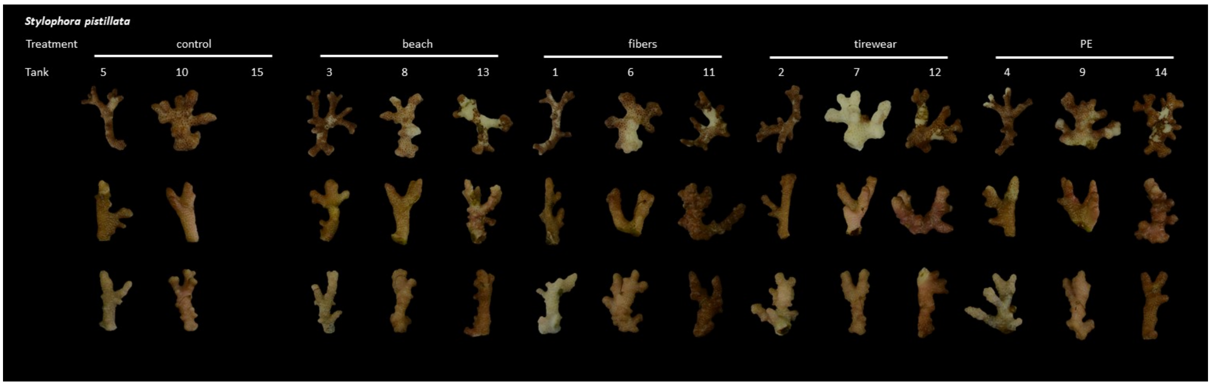


**Supplementary Figure 32.** Overview of all fragments of the stony coral species *Stylophora pistillata* after 10 weeks of combined microplastic and heat stress treatment. The displayed pictures were used to calculate brightness as a measure of coral bleaching. Fragments from tank 15 are missing due to a technical error in the aquarium.

### Supplementary tables

**Supplementary Table 1.** Time points of independent experiments on the combined effects of microplastic pollution and heat stress on stony coral species.

| | Addition of microplastic particles | Heat ramping phase | End of experiment |
|---|---|---|---|
| Experiment 1 | 01.10.2019 | 07.-14.11.2019 | 29.11.2019 |
| Experiment 2 | 12.09.2018 | 11.-26.10.2018 | 23.11.2018 |
| Experiment 3 | 19.06.2019 | 21.-27.08.2019 | 27.08.2019 |

**Supplementary Table 2.** Time points of measurements of photosynthetic efficiency of stony coral species in independent experiments on the combined effects of microplastic pollution and heat stress.

| Experimental phase | Experiment 1 | Experiment 2 | Experiment 3 |
|---|---|---|---|
| no treatment | 23.-26.09.2019 | 11.09.2018 | |
| microplastic | 21.-28.10.2019 | 18.09.2018 | |
| microplastic | | 25.09.2018 | |
| microplastic | | 02.10.2018 | |
| microplastic | | 09.10.2018 | |
| microplastic + heat | 14.-26.11.2019 | 16.10.2018 | 22.-27.08.2019 |
| microplastic + heat | | 23.10.2018 | |
| microplastic + heat | | 27.10.2018 | |
| microplastic + heat | | 31.10.2018 | |

**Supplementary Table 3.** Time points for standardized photos to assess tissue brightness and bleaching of stony coral species in independent experiments on the combined effects of microplastic pollution and heat stress.

| Experimental phase | Experiment 1 | Experiment 2 | Experiment 3 |
|---|---|---|---|
| no treatment | 27.09.2019 | | |
| microplastic | 30.10.2019 | 11.10.2018 | |
| microplastic + heat | 16.-26.11.2019 | 18.10.2018 | 22.-27.08.2019 |
| microplastic + heat | | 25.10.2018 | |
| microplastic + heat | | 01.11.2018 | |

**Supplementary Table 4.** Information on coral colony origin, collection, and permit numbers.

| Species | Colony | Origin | Collection | Permit number (CITES) |
|---|---|---|---|---|
| *A. muricata* | A-C | Indonesia | 12.2007 | 14846/IV/SATS-LN/2007 |
| *M. digitata* | A-C | Indonesia | 10.2014 | Transfer agreement with Customs authority (no. L11/15) |
| *P. cylindrica* | A-C | Indonesia | 2017 | 17nl241924/11 |
| *P. verrucosa* | A & C, D | Indonesia | 12.2007 | 14846/IV/SATS-LN/2007 |
| *P. verrucosa* | B | Indonesia | 04.2014 | 14NL214371/11 |
| *P. verrucosa* | E-J | Saudi Arabia | 03.2019 | 19-SA-000091-PD |
| *S. pistillata* | A-D | Fiji | 2016 | 15NL229192/11 |
| *S. pistillata* | E-J | Saudi Arabia | 03.2019 | 19-SA-000092-PD |

**Supplementary Table 5**. Results of statistical analyses of Experiment 1, assessing the effects of heat stress (heat), microplastic exposure (plastic), and its interaction (interaction) on physiological parameters (maximum photosynthetic efficiency (Fv/Fm), effective photosynthetic efficiency (ΔF/Fm'), bleaching, and mortality) of *Pocillopora verrucosa* and *Stylophora pistillata*. Overall impacts of heat stress and microplastic exposure on Fv/Fm, ΔF/Fm', and bleaching are analyzed with linear mixed-effects models (lmer) on all data. Impacts of microplastic under ambient temperature and under heat stress are analyzed separately with linear mixed-effects models on sub-datasets. Overall impacts on mortality are derived from log-rank statistics (heat stress) and cox models (microplastic exposure) on all data. Heat (ambient temperature vs. heat stress), plastic (control vs. microplastic), or treatment (concentrations: control, c2.5, c25, c250, c2500) were set as fixed factors and colony (col), origin and nubbin identity (ID) were set as random factors if applicable. Number of observations (n), estimates, standard errors, z-values, and p-values are given with model specifications. Bold values indicate significance (P<0.05).

| Response variable | Species | dataset | n | Factor | Estimate | Std. Error | z-value | p-value | Model specifications |
|---|---|---|---|---|---|---|---|---|---|
| Maximum photosynthetic efficiency (Fv/Fm) | *P. verrucosa* | all data | 240 | heat | -0.2479 | 0.0261 | -9.5100 | **<0.0001** | lmer, YII^2 ~ heat * plastic + (1 \| col) + (1 \| origin) + (1 \| ID) |
| | | | | plastic | -0.0070 | 0.0206 | -0.3400 | 0.7340 | |
| | | | | interaction | 0.0234 | 0.0292 | 0.8020 | 0.4220 | |
| | | ambient temp | 120 | plastic | -0.0042 | 0.0041 | -1.0260 | 0.3050 | lmer, YII ~ plastic + (1 \| col) + (1 \| origin) |
| | | | | control - c2.5 | 0.0078 | 0.0052 | 1.5050 | 0.5290 | lmer, YII ~ treat + (1 \| col) + (1 \| origin) |
| | | | | control - c25 | 0.0035 | 0.0052 | 0.6810 | 1.0000 | |
| | | | | control - c250 | 0.0016 | 0.0053 | 0.3110 | 1.0000 | |
| | | | | control - c2500 | 0.0037 | 0.0052 | 0.7050 | 1.0000 | |
| | | heat stress | 120 | plastic | 0.0260 | 0.0254 | 1.0250 | 0.3055 | lmer, YII ~ plastic + (1 \| col) + (1 \| origin) |
| | | | | control - c2.5 | -0.0434 | 0.0314 | -1.3810 | 0.6690 | lmer, YII ~ treat + (1 \| col) + (1 \| origin) |
| | | | | control - c25 | 0.0271 | 0.0314 | 0.8630 | 1.0000 | |
| | | | | control - c250 | -0.0410 | 0.0314 | -1.3060 | 0.7660 | |

| | | | | | | | | | |
|---|---|---|---|---|---|---|---|---|---|
| | | | | control - c2500 | -0.0466 | 0.0314 | -1.4840 | 0.5520 | |
| | *S. pistillata* | all data | 240 | heat | -0.2893 | 0.0213 | -13.5590 | **<0.0001** | lmer, YII^2 ~ heat * plastic + (1 \| col) + (1 \| origin) + (1 \| ID) |
| | | | | plastic | -0.0027 | 0.0169 | -0.1600 | 0.8730 | |
| | | | | interaction | 0.0489 | 0.0239 | 2.0520 | **0.0402** | |
| | | ambient temp | 120 | plastic | -0.0021 | 0.0056 | -0.3740 | 0.7090 | lmer, YII ~ plastic + (1 \| col) + (1 \| origin) |
| | | | | control - c2.5 | 0.0020 | 0.0072 | 0.2720 | 1.0000 | lmer, YII ~ treat + (1 \| col) + (1 \| origin) |
| | | | | control - c25 | 0.0015 | 0.0072 | 0.2020 | 1.0000 | |
| | | | | control - c250 | 0.0046 | 0.0072 | 0.6360 | 1.0000 | |
| | | | | control - c2500 | 0.0004 | 0.0072 | 0.0580 | 1.0000 | |
| | | heat stress | 120 | plastic | 0.0462 | 0.0173 | 2.6730 | **0.0075** | lmer, YII ~ plastic + (1 \| col) + (1 \| origin) |
| | | | | control - c2.5 | -0.0521 | 0.0217 | -2.4020 | 0.0653 | lmer, YII ~ treat + (1 \| col) + (1 \| origin) |
| | | | | control - c25 | -0.0211 | 0.0217 | -0.9740 | 1.0000 | |
| | | | | control - c250 | -0.0436 | 0.0217 | -2.0090 | 0.1783 | |
| | | | | control - c2500 | -0.0681 | 0.0217 | -3.1360 | **0.0069** | |
| Effective photosynthetic efficiency (ΔF/Fm') | *P. verrucosa* | all data | 720 | heat | -0.2156 | 0.0133 | -16.1820 | **<0.0001** | lmer, YII^2 ~ heat * plastic + (1 \| col) + (1 \| origin) + (1 \| ID) |
| | | | | plastic | -0.0010 | 0.0105 | -0.0970 | 0.9225 | |
| | | | | interaction | 0.0303 | 0.0149 | 2.0350 | **0.0418** | |
| | | ambient temp | 360 | plastic | 0.0010 | 0.0028 | -0.3630 | 0.7160 | lmer, YII ~ plastic + (1 \| col) + (1 \| origin) |
| | | | | control - c2.5 | 0.0067 | 0.0035 | 1.9260 | 0.2160 | lmer, YII ~ treat + (1 \| col) + (1 \| origin) |
| | | | | control - c25 | -0.0015 | 0.0035 | -0.4420 | 1.0000 | |
| | | | | control - c250 | 0.0004 | 0.0035 | 0.1100 | 1.0000 | |
| | | | | control - c2500 | -0.0015 | 0.0035 | -0.4220 | 1.0000 | |
| | | heat stress | 360 | plastic | 0.0364 | 0.0181 | 2.0060 | **0.0448** | lmer, YII ~ plastic + (1 \| col) + (1 \| origin) |
| | | | | control - c2.5 | -0.0383 | 0.0221 | -1.7310 | 0.3338 | lmer, YII ~ treat + (1 \| col) + (1 \| origin) |
| | | | | control - c25 | 0.0064 | 0.0221 | 0.2920 | 1.0000 | |
| | | | | control - c250 | -0.0503 | 0.0221 | -2.2750 | 0.0917 | |
| | | | | control - c2500 | -0.0633 | 0.0221 | -2.8630 | **0.0168** | |
| | *S. pistillata* | all data | 720 | heat | -0.1894 | 0.0110 | -17.2510 | **<0.0001** | lmer, YII^2 ~ heat * plastic + (1 \| col) + (1 \| origin) + (1 \| ID) |
| | | | | plastic | 0.0029 | 0.0097 | 0.2960 | 0.7674 | |
| | | | | interaction | 0.0291 | 0.0123 | 2.3690 | **0.0178** | |
| | | ambient temp | 360 | plastic | 0.0025 | 0.0022 | 1.1420 | 0.2530 | lmer, YII ~ plastic + (1 \| col) + (1 \| origin) |

| | | | | | | | | | |
|---|---|---|---|---|---|---|---|---|---|
| | | | | control - c2.5 | 0.0031 | 0.0027 | 1.1540 | 0.9936 | lmer, YII ~ treat + (1 \| col) + (1 \| origin) |
| | | | | control - c25 | -0.0029 | 0.0027 | -1.0890 | 1.0000 | |
| | | | | control - c250 | -0.0036 | 0.0027 | -1.3450 | 0.7148 | |
| | | | | control - c2500 | -0.0067 | 0.0027 | -2.5080 | **0.0485** | |
| | | heat stress | 360 | plastic | 0.0319 | 0.0144 | 2.2150 | **0.0268** | lmer, (YII)^2 ~ plastic + (1 \| col) + (1 \| origin) |
| | | | | control - c2.5 | -0.0393 | 0.0176 | -2.2340 | 0.1020 | lmer, (YII)^2 ~ treat + (1 \| col) + (1 \| origin) |
| | | | | control - c25 | -0.0082 | 0.0176 | -0.4680 | 1.0000 | |
| | | | | control - c250 | -0.0179 | 0.0176 | -1.0160 | 1.0000 | |
| | | | | control - c2500 | -0.0623 | 0.0176 | -3.5410 | **0.0016** | |
| Bleaching (tissue Brightness) | *P. verrucosa* | all data | 240 | heat | 55.0920 | 5.9690 | 9.2290 | **<0.0001** | lmer, gray ~ heat * plastic + (1 \| col) + (1 \| origin) + (1 \| ID) |
| | | | | plastic | 8.4740 | 4.7190 | 1.7960 | 0.0726 | |
| | | | | interaction | -12.4100 | 6.6740 | -1.8600 | 0.0630 | |
| | | ambient temp | 120 | plastic | 8.4740 | 3.8410 | 2.2060 | **0.0274** | lmer, gray ~ plastic + (1 \| col) + (1 \| origin) |
| | | | | control - c2.5 | -7.5340 | 4.8360 | -1.5580 | 0.4771 | lmer, gray ~ treat + (1 \| col) + (1 \| origin) |
| | | | | control - c25 | -8.3010 | 4.8360 | -1.7160 | 0.3444 | |
| | | | | control - c250 | -4.2810 | 4.8360 | -0.8850 | 1.0000 | |
| | | | | control - c2500 | -13.7780 | 4.8360 | -2.8490 | **0.0175** | |
| | | heat stress | 120 | plastic | -3.9370 | 3.6840 | -1.0690 | 0.2850 | lmer, gray ~ plastic + (1 \| col) + (1 \| origin) |
| | | | | control - c2.5 | 6.8550 | 4.6500 | 1.4740 | 0.5620 | lmer, gray ~ treat + (1 \| col) + (1 \| origin) |
| | | | | control - c25 | -1.1600 | 4.6500 | -0.2490 | 1.0000 | |
| | | | | control - c250 | 5.7830 | 4.6500 | 1.2440 | 0.8540 | |
| | | | | control - c2500 | 4.2690 | 4.6500 | 0.9180 | 1.0000 | |
| | *S. pistillata* | all data | 240 | heat | 61.2080 | 5.3990 | 11.3380 | **<0.0001** | lmer, gray ~ heat * plastic + (1 \| col) + (1 \| origin) + (1 \| ID) |
| | | | | plastic | -7.1040 | 4.2680 | -1.6650 | 0.0960 | |
| | | | | interaction | -11.9250 | 6.0360 | -1.9760 | **0.0482** | |
| | | ambient temp | 120 | plastic | -7.1040 | 3.4590 | -2.0540 | **0.0400** | lmer, gray ~ plastic + (1 \| col) + (1 \| origin) |
| | | | | control - c2.5 | 8.4960 | 4.4210 | 1.9220 | 0.2180 | lmer, gray ~ treat + (1 \| col) + (1 \| origin) |

| | | | | | | | | | |
|---|---|---|---|---|---|---|---|---|---|
| | | | | control - c25 | 5.3000 | 4.4210 | 1.1990 | 0.9220 | |
| | | | | control - c250 | 8.2500 | 4.4210 | 1.8660 | 0.2480 | |
| | | | | control - c2500 | 6.3700 | 4.4210 | 1.4410 | 0.5990 | |
| | | heat stress | 120 | plastic | -19.0290 | 4.6260 | -4.1140 | **<0.0001** | lmer, gray ~ plastic + (1 \| col) + (1 \| origin) |
| | | | | control - c2.5 | 17.0050 | 5.8780 | 2.8930 | **0.0153** | lmer, gray ~ treat + (1 \| col) + (1 \| origin) |
| | | | | control - c25 | 15.4320 | 5.8780 | 2.6250 | **0.0346** | |
| | | | | control - c250 | 22.6430 | 5.8780 | 3.8520 | **0.0005** | |
| | | | | control - c2500 | 21.0370 | 5.8780 | 3.5790 | **0.0014** | |
| Mortality | *P. verrucosa* | all data | | heat | NA | NA | 14.7020 | **<0.0001** | logrank, Surv(time, mort) by heat |
| | | | 120 | plastic | -0.3621 | 0.2323 | -1.5590 | 0.1190 | coxme, (time,mort)~plastic + (1\|col) + (1\|origin) |
| | | | 120 | control - c2.5 | 0.3780 | 0.2970 | 1.2720 | 0.5238 | coxme, (time,mort)~treat + (1\|col) + (1\|origin) |
| | | | | control - c25 | -0.2786 | 0.3009 | -0.9260 | 0.7681 | |
| | | | | control - c250 | 0.1134 | 0.2948 | 0.3850 | 0.9870 | |
| | | | | control - c2500 | 0.9409 | 0.3070 | 3.0640 | **0.0082** | |
| | *S. pistillata* | all data | | heat | NA | NA | 11.2480 | **<0.0001** | logrank, Surv(time, mort) by heat |
| | | | 120 | plastic | -1.0943 | 0.2856 | -3.8320 | **0.0001** | coxme, (time,mort)~plastic + (1\|col) + (1\|origin) |
| | | | 120 | control - c2.5 | 1.0575 | 0.3620 | 2.9210 | **0.0139** | coxme, (time,mort)~treat + (1\|col) + (1\|origin) |
| | | | | control - c25 | 1.1019 | 0.3568 | 3.0880 | **0.0081** | |
| | | | | control - c250 | 1.0403 | 0.3597 | 2.8920 | **0.0153** | |
| | | | | control - c2500 | 1.2882 | 0.3714 | 3.4680 | **0.0021** | |

**Supplementary Table 6**. Results of statistical analyses of Experiment 2, assessing the effects of heat stress (heat), microplastic exposure (plastic), and its interaction (interaction) on physiological parameters (maximum photosynthetic efficiency (Fv/Fm), effective photosynthetic efficiency (ΔF/Fm'), bleaching, and tissue necrosis) of *Acropora muricata, Montipora digitata*, *Porites cylindrica*, *Pocillopora verrucosa*, and *Stylophora pistillata*. Overall impacts of heat stress and microplastic exposure are analyzed with linear mixed-effects models (lmer) on all data. Impacts of microplastic under ambient temperature and under heat stress are analyzed separately with linear mixed-effects models on sub-datasets. Heat (ambient temperature vs. heat stress), or plastic (control vs. microplastic) were set as fixed factors and colony (col) and nubbin identity (ID) were set as random factors if applicable. Number of observations* (n), estimates, standard errors, z-values, and p-values are given with model specifications. Bold values indicate significance (P<0.05).

* number of observations for effective photosynthetic efficiency might be lower when a repeated measurement on the same nubbin was not possible due to nubbin size or status; no observations for tissue brightness of colony Pve_C at the last timepoint as the nubbins died before pictures could be taken; missing datapoint in tissue necrosis for Spi_A_1_2 due to corrupt 3D model.

<table>
<tr><th>Response variable</th><th>Species</th><th>Dataset</th><th>n</th><th>Factor</th><th>Estimate</th><th>Std. Error</th><th>z-value</th><th>p-value</th><th>model specifications</th></tr>
<tr><td>Maximum photosynthetic efficiency (Fv/Fm)</td><td>A. muricata</td><td>all data</td><td>36</td><td>heat</td><td>-0.2039</td><td>0.0646</td><td>-3.1560</td><td>0.0016</td><td>lmer, YII ~ heat * plastic + (1 | col) + (1 | ID)</td></tr>
<tr><td></td><td></td><td></td><td></td><td>plastic</td><td>0.0359</td><td>0.0646</td><td>0.5560</td><td>0.5785</td><td></td></tr>
<tr><td></td><td></td><td></td><td></td><td>interaction</td><td>-0.0263</td><td>0.0914</td><td>-0.2880</td><td>0.7732</td><td></td></tr>
<tr><td></td><td></td><td>ambient temp</td><td>18</td><td>plastic</td><td>0.0359</td><td>0.0416</td><td>0.8630</td><td>0.3880</td><td>lmer, YII ~ plastic + (1 | col)</td></tr>
<tr><td></td><td></td><td>heat stress</td><td>18</td><td>plastic</td><td>0.0096</td><td>0.0709</td><td>0.1350</td><td>0.8928</td><td>lmer, YII ~ plastic + (1 | col)</td></tr>
<tr><td></td><td>M. digitata</td><td>all data</td><td>36</td><td>heat</td><td>-0.2463</td><td>0.0417</td><td>-5.9030</td><td><0.0001</td><td>lmer, YII ~ heat * plastic + (1 | col) + (1 | ID)</td></tr>
<tr><td></td><td></td><td></td><td></td><td>plastic</td><td>-0.0356</td><td>0.0548</td><td>-0.6490</td><td>0.5160</td><td></td></tr>
<tr><td></td><td></td><td></td><td></td><td>interaction</td><td>0.0659</td><td>0.0590</td><td>1.1160</td><td>0.2640</td><td></td></tr>
</table>

| | | | | | | | | | |
|---|---|---|---|---|---|---|---|---|---|
| | | ambient temp | 18 | plastic | -0.0356 | 0.0426 | -0.8360 | 0.4030 | lmer, YII ~ plastic + (1 \| col) |
| | | heat stress | 18 | plastic | 0.0303 | 0.0624 | 0.4860 | 0.6269 | lmer, YII ~ plastic + (1 \| col) |
| | *P. cylindrica* | all data | 36 | heat | -0.2693 | 0.0403 | -6.6770 | **<0.0001** | lmer, YII ~ heat * plastic + (1 \| col) + (1 \| ID) |
| | | | | plastic | 0.0101 | 0.0403 | 0.2510 | 0.8020 | |
| | | | | interaction | 0.0014 | 0.0570 | 0.0250 | 0.9800 | |
| | | ambient temp | 18 | plastic | 0.0101 | 0.0343 | 0.2940 | 0.7680 | lmer, YII ~ plastic + (1 \| col) |
| | | heat stress | 18 | plastic | 0.0116 | 0.0447 | 0.2580 | 0.7960 | lmer, YII ~ plastic + (1 \| col) |
| | *P. verrucosa* | all data | 36 | heat | -0.3452 | 0.0458 | -7.5400 | **<0.0001** | lmer, YII ~ heat * plastic + (1 \| col) + (1 \| ID) |
| | | | | plastic | -0.0904 | 0.0458 | -1.9750 | **0.0482** | |
| | | | | interaction | -0.0114 | 0.0648 | -0.1770 | 0.8597 | |
| | | ambient temp | 18 | plastic | -0.0904 | 0.0331 | -2.7330 | **0.0063** | lmer, YII ~ plastic + (1 \| col) |
| | | heat stress | 18 | plastic | -0.1019 | 0.0488 | -2.0860 | **0.0370** | lmer, YII ~ plastic + (1 \| col) |
| | *S. pistillata* | all data | 36 | heat | -0.2647 | 0.0422 | -6.2720 | **<0.0001** | lmer, YII ~ heat * plastic + (1 \| col) + (1 \| ID) |
| | | | | plastic | 0.0049 | 0.0422 | 0.1160 | 0.9080 | |
| | | | | interaction | -0.0719 | 0.0597 | -1.2050 | 0.2280 | |
| | | ambient temp | 18 | plastic | 0.0049 | 0.0383 | 0.1280 | 0.8980 | lmer, YII ~ plastic + (1 \| col) |
| | | heat stress | 18 | plastic | -0.0670 | 0.0442 | -1.5150 | 0.1300 | lmer, YII ~ plastic + (1 \| col) |
| Effective photosynthetic efficiency (ΔF/Fm') | *A. muricata* | all data | 108 | heat | -0.1124 | 0.0279 | -4.0220 | **0.0001** | lmer, YII ~ heat * plastic + (1 \| col) + (1 \| ID) |
| | | | | plastic | 0.0053 | 0.0466 | 0.1140 | 0.9090 | |
| | | | | interaction | -0.0273 | 0.0395 | -0.6910 | 0.4900 | |
| | | ambient temp | 54 | plastic | 0.0053 | 0.0254 | 0.2100 | 0.8340 | lmer, YII ~ plastic + (1 \| col) |
| | | heat stress | 54 | plastic | -0.0220 | 0.0721 | -0.3050 | 0.7610 | lmer, YII ~ plastic + (1 \| col) |
| | *M. digitata* | all data | 107 | heat | -0.0311 | 0.0279 | -1.1130 | 0.2660 | lmer, YII ~ heat * plastic + (1 \| col) + (1 \| ID) |
| | | | | plastic | -0.0260 | 0.0301 | -0.8630 | 0.3880 | |
| | | | | interaction | 0.0130 | 0.0393 | 0.3300 | 0.7410 | |
| | | ambient temp | 54 | plastic | -0.0260 | 0.0331 | -0.7840 | 0.4330 | lmer, YII ~ plastic + (1 \| col) |
| | | heat stress | 53 | plastic | -0.0124 | 0.0354 | -0.3500 | 0.7260 | lmer, YII ~ plastic + (1 \| col) |
| | *P. cylindrica* | all data | 108 | heat | -0.0175 | 0.0242 | -0.7240 | 0.4690 | lmer, YII ~ heat * plastic + (1 \| col) + (1 \| ID) |
| | | | | plastic | 0.0223 | 0.0274 | 0.8120 | 0.4170 | |
| | | | | interaction | -0.0049 | 0.0342 | -0.1430 | 0.8860 | |
| | | ambient temp | 54 | plastic | 0.0223 | 0.0246 | 0.9040 | 0.3660 | lmer, YII ~ plastic + (1 \| col) |
| | | heat stress | 54 | plastic | 0.0174 | 0.0271 | 0.6420 | 0.5210 | lmer, YII ~ plastic + (1 \| col) |
| | *P. verrucosa* | all data | 106 | heat | -0.1184 | 0.0254 | -4.6530 | **<0.0001** | lmer, YII ~ heat * plastic + (1 \| col) + (1 \| ID) |

| | | | | | | | | | |
|---|---|---|---|---|---|---|---|---|---|
| | | | | plastic | -0.0104 | 0.0274 | -0.3790 | 0.7050 | |
| | | | | interaction | 0.0058 | 0.0360 | 0.1610 | 0.8720 | |
| | | ambient temp | 54 | plastic | -0.0104 | 0.0240 | -0.4330 | 0.6650 | lmer, YII ~ plastic + (1 \| col) |
| | | heat stress | 52 | plastic | -0.0055 | 0.0350 | -0.1570 | 0.8750 | lmer, YII ~ plastic + (1 \| col) |
| | *S. pistillata* | all data | 107 | heat | -0.0504 | 0.0265 | -1.9000 | 0.0575 | lmer, YII ~ heat * plastic + (1 \| col) + (1 \| ID) |
| | | | | plastic | -0.0052 | 0.0337 | -0.1530 | 0.8782 | |
| | | | | interaction | -0.0664 | 0.0377 | -1.7610 | 0.0782 | |
| | | ambient temp | 53 | plastic | -0.0054 | 0.0221 | -0.2440 | 0.8070 | lmer, YII ~ plastic + (1 \| col) |
| | | heat stress | 54 | plastic | -0.0716 | 0.0428 | -1.6740 | 0.0941 | lmer, YII ~ plastic + (1 \| col) |
| Bleaching (tissue brightness) | *A. muricata* | all data | 36 | heat | 121.1540 | 5.9830 | 20.2490 | **<0.0001** | lmer, gray ~ heat * plastic + (1 \| col) + (1 \| ID) |
| | | | | plastic | 5.2670 | 7.3910 | 0.7130 | 0.4760 | |
| | | | | interaction | -4.6550 | 8.4610 | -0.5500 | 0.5820 | |
| | | ambient temp | 18 | plastic | 5.2670 | 8.9480 | 0.5890 | 0.5560 | lmer, gray ~ plastic + (1 \| col) |
| | | heat stress | 18 | plastic | 0.6118 | 4.6743 | 0.1310 | 0.8960 | lmer, gray ~ plastic + (1 \| col) |
| | *M. digitata* | all data | 36 | heat | 35.8400 | 5.1000 | 7.0270 | **<0.0001** | lmer, gray ~ heat * plastic + (1 \| col) + (1 \| ID) |
| | | | | plastic | -22.0530 | 7.7670 | -2.8390 | **0.0045** | |
| | | | | interaction | 3.8400 | 7.2130 | 0.5320 | 0.5945 | |
| | | ambient temp | 18 | plastic | -22.0530 | 7.0190 | -3.1420 | **0.0017** | lmer, gray ~ plastic + (1 \| col) |
| | | heat stress | 18 | plastic | -18.2130 | 7.9980 | -2.2770 | **0.0228** | lmer, gray ~ plastic + (1 \| col) |
| | *P. cylindrica* | all data | 36 | heat | 45.1986 | 3.8470 | 11.7490 | **<0.0001** | lmer, gray ~ heat * plastic + (1 \| col) + (1 \| ID) |
| | | | | plastic | -11.4440 | 7.2642 | -1.5750 | 0.1150 | |
| | | | | interaction | 0.5946 | 5.4404 | 0.1090 | 0.9130 | |
| | | ambient temp | 18 | plastic | -11.4440 | 7.1410 | -1.6020 | 0.1090 | lmer, gray ~ plastic + (1 \| col) |
| | | heat stress | 18 | plastic | -10.8490 | 7.4280 | -1.4610 | 0.1440 | lmer, gray ~ plastic + (1 \| col) |
| | *P. verrucosa* | all data | 30 | heat | 107.4530 | 7.5920 | 14.1540 | **<0.0001** | lmer, gray ~ heat * plastic + (1 \| col) + (1 \| ID) |
| | | | | plastic | 23.2810 | 7.3590 | 3.1640 | **0.0016** | |
| | | | | interaction | -15.3810 | 10.4080 | -1.4780 | 0.1395 | |
| | | ambient temp | 18 | plastic | 23.2800 | 9.1500 | 2.5440 | **0.0109** | lmer, gray ~ plastic + (1 \| col) |
| | | heat stress | 12 | plastic | 9.5860 | 3.3990 | 2.8210 | **0.0048** | lmer, gray ~ plastic + (1 \| col) |
| | *S. pistillata* | all data | 36 | heat | 46.0230 | 4.3610 | 10.5540 | **<0.0001** | lmer, gray ~ heat * plastic + (1 \| col) + (1 \| ID) |
| | | | | plastic | 1.4490 | 11.7740 | 0.1230 | 0.9020 | |
| | | | | interaction | 1.3660 | 6.1670 | 0.2220 | 0.8250 | |
| | | ambient temp | 18 | plastic | 1.4490 | 11.0180 | 0.1320 | 0.8950 | lmer, gray ~ plastic + (1 \| col) |
| | | heat stress | 18 | plastic | 2.8150 | 12.0990 | 0.2330 | 0.8160 | lmer, gray ~ plastic + (1 \| col) |

| | | | | | | | | | |
|---|---|---|---|---|---|---|---|---|---|
| Tissue necrosis (percent healthy tissue) | *A. muricata* | all data | 36 | heat | -4.1808 | 1.8142 | -2.3040 | **0.0212** | lmer, healthy ~ heat * plastic + (1 \| col) + (1 \| ID) |
| | | | | plastic | -0.2178 | 1.8142 | -0.1200 | 0.9044 | |
| | | | | interaction | -1.6392 | 2.5657 | -0.6390 | 0.5229 | |
| | | ambient temp | 18 | plastic | -0.2178 | 0.2178 | -1.0000 | 0.3170 | lmer, healthy ~ plastic + (1 \| col) |
| | | heat stress | 18 | plastic | -1.8570 | 2.1190 | -0.8760 | 0.3810 | lmer, healthy ~ plastic + (1 \| col) |
| | *M. digitata* | all data | 36 | heat | -57.7950 | 9.4420 | -6.1210 | **<0.0001** | lmer, healthy ~ heat * plastic + (1 \| col) + (1 \| ID) |
| | | | | plastic | 1.3750 | 9.4420 | 0.1460 | 0.8840 | |
| | | | | interaction | 6.8850 | 13.3540 | 0.5160 | 0.6060 | |
| | | ambient temp | 18 | plastic | 1.3751 | 0.7139 | 1.9260 | 0.0541 | lmer, healthy ~ plastic + (1 \| col) |
| | | heat stress | 18 | plastic | 8.2600 | 2.0490 | 4.0310 | **0.0001** | lmer, healthy ~ plastic + (1 \| col) |
| | *P. cylindrica* | all data | 36 | heat | -32.2190 | 9.4470 | -3.4110 | **0.0006** | lmer, healthy ~ heat * plastic + (1 \| col) + (1 \| ID) |
| | | | | plastic | 1.8540 | 9.4470 | 0.1960 | 0.8444 | |
| | | | | interaction | -9.4210 | 13.3600 | -0.7050 | 0.4807 | |
| | | ambient temp | 18 | plastic | 1.8540 | 1.7220 | 1.0770 | 0.2820 | lmer, healthy ~ plastic + (1 \| col) |
| | | heat stress | 18 | plastic | -7.5680 | 10.1360 | -0.7470 | 0.4553 | lmer, healthy ~ plastic + (1 \| col) |
| | *P. verrucosa* | all data | 36 | heat | -61.1050 | 10.3397 | -5.9100 | **<0.0001** | lmer, healthy ~ heat * plastic + (1 \| col) + (1 \| ID) |
| | | | | plastic | 0.0201 | 10.3397 | 0.0020 | 0.9980 | |
| | | | | interaction | -8.8728 | 14.6226 | -0.6070 | 0.5440 | |
| | | ambient temp | 18 | plastic | 0.0201 | 0.2985 | 0.0670 | 0.9460 | lmer, healthy ~ plastic + (1 \| col) |
| | | heat stress | 18 | plastic | -8.8530 | 9.9760 | -0.8870 | 0.3748 | lmer, healthy ~ plastic + (1 \| col) |
| | *S. pistillata* | all data | 35 | heat | -66.8500 | 11.3300 | -5.8990 | **<0.0001** | lmer, healthy ~ heat * plastic + (1 \| col) + (1 \| ID) |
| | | | | plastic | 0.0000 | 11.3300 | 0.0000 | 1.0000 | |
| | | | | interaction | 1.0210 | 16.2900 | 0.0630 | 0.9500 | |
| | | ambient temp | 18 | plastic | 0.0000 | <0.0001 | 0.0000 | 1.0000 | lmer, healthy ~ plastic + (1 \| col) |
| | | heat stress | 17 | plastic | -1.0290 | 3.1750 | -0.3240 | 0.7460 | lmer, healthy ~ plastic + (1 \| col) |

**Supplementary Table 7**. Results of statistical analyses of Experiment 3 assessing the effects of different microplastic mixtures (fibers, tirewear, beach and PE) on physiological parameters (maximum photosynthetic efficiency (Fv/Fm), effective photosynthetic efficiency (ΔF/Fm'), bleaching, and tissue necrosis) of *Pocillopora verrucosa* and *Stylophora pistillata* under heat stress. Impacts of exposure to microplastic mixtures under heat stress are derived from linear mixed-effects models (lmer) on all data. Treatment (control, fibers, tirewear, beach and PE) was set as fixed factor and colony (col), origin, and nubbin identity (ID) were set as random factors if applicable. Number of observations* (n), estimates, standard errors, z-values, and p-values are given with model specifications. Bold values indicate significance ($P<0.05$).

* number of observations for effective photosynthetic efficiency might be lower as all measurements from tank 15 were excluded after a temperature control error during the course of the experiment (22.08.2019)

<table>
<tr><th>Response variable</th><th>Species</th><th>n</th><th>Comparison</th><th>Estimate</th><th>Std. Error</th><th>z-value</th><th>p-value</th><th>model specifications</th></tr>
<tr><td>Maximum photosynthetic efficiency (Fv/Fm)</td><td>P. verrucosa</td><td>43</td><td>control - fibers</td><td>-0.0011</td><td>0.0218</td><td>-0.0490</td><td>1.0000</td><td>lmer, YII ~ treat + (1 | col) + (1 | origin)</td></tr>
<tr><td></td><td></td><td></td><td>control - tirewear</td><td>-0.0104</td><td>0.0218</td><td>-0.4770</td><td>1.0000</td><td></td></tr>
<tr><td></td><td></td><td></td><td>control - beach</td><td>0.0093</td><td>0.0218</td><td>0.4240</td><td>1.0000</td><td></td></tr>
<tr><td></td><td></td><td></td><td>control - PE</td><td>-0.0007</td><td>0.0218</td><td>-0.0340</td><td>1.0000</td><td></td></tr>
<tr><td></td><td>S. pistillata</td><td>42</td><td>control - fibers</td><td>0.0082</td><td>0.0247</td><td>0.3330</td><td>1.0000</td><td>lmer, YII ~ treat + (1 | col) + (1 | origin)</td></tr>
<tr><td></td><td></td><td></td><td>control - tirewear</td><td>0.0118</td><td>0.0247</td><td>0.4780</td><td>1.0000</td><td></td></tr>
<tr><td></td><td></td><td></td><td>control - beach</td><td>0.0260</td><td>0.0247</td><td>1.0540</td><td>1.0000</td><td></td></tr>
<tr><td></td><td></td><td></td><td>control - PE</td><td>0.0074</td><td>0.0247</td><td>0.3000</td><td>1.0000</td><td></td></tr>
<tr><td>Effective photosynthetic efficiency (ΔF/Fm')</td><td>P. verrucosa</td><td>56</td><td>control - fibers</td><td>0.0575</td><td>0.0181</td><td>3.1760</td><td>0.0060</td><td>lmer, YII ~ treat + (1 | col) + (1 | origin) + (1|ID)</td></tr>
<tr><td></td><td></td><td></td><td>control - tirewear</td><td>0.0330</td><td>0.0181</td><td>1.8250</td><td>0.2720</td><td></td></tr>
<tr><td></td><td></td><td></td><td>control - beach</td><td>0.0375</td><td>0.0181</td><td>2.0720</td><td>0.1531</td><td></td></tr>
</table>

| | | | | | | | |
|---|---|---|---|---|---|---|---|
| | | | control - PE | 0.0167 | 0.0181 | 0.9240 | 1.0000 | |
| | *S. pistillata* | 74 | control - fibers | 0.0135 | 0.0336 | 0.4030 | 1.0000 | lmer, YII ~ treat + (1 \| col) + (1 \| origin) + (1\|ID) |
| | | | control - tirewear | 0.0447 | 0.0336 | 1.3330 | 0.7300 | |
| | | | control - beach | 0.0487 | 0.0336 | 1.4520 | 0.5860 | |
| | | | control - PE | -0.0025 | 0.0336 | -0.0760 | 1.0000 | |
| Bleaching (tissue brightness) | *P. verrucosa* | 43 | control - fibers | 1.7070 | 10.6510 | 0.1600 | 1.0000 | lmer, gray ~ treat + (1 \| col) + (1 \| origin) |
| | | | control - tirewear | 1.0860 | 10.6510 | 0.1020 | 1.0000 | |
| | | | control - beach | -7.4210 | 10.6510 | -0.6970 | 1.0000 | |
| | | | control - PE | -8.0380 | 10.6510 | -0.7550 | 1.0000 | |
| | *S. pistillata* | 42 | control - fibers | 0.0371 | 0.0646 | 0.5740 | 1.0000 | lmer, gray ~ treat + (1 \| col) + (1 \| origin) |
| | | | control - tirewear | -0.0473 | 0.0646 | -0.7320 | 1.0000 | |
| | | | control - beach | -0.0400 | 0.0646 | -0.6190 | 1.0000 | |
| | | | control - PE | 0.0463 | 0.0646 | 0.7160 | 1.0000 | |
| Tissue necrosis (percent healthy tissue) | *P. verrucosa* | 43 | control - fibers | -4.8260 | 11.7670 | -0.4100 | 1.0000 | lmer, healthy ~ treat + (1 \| col) + (1 \| origin) |
| | | | control - tirewear | -14.2400 | 11.7670 | -1.2100 | 0.9050 | |
| | | | control - beach | 1.7250 | 11.7670 | 0.1470 | 1.0000 | |
| | | | control - PE | -7.6550 | 11.7670 | -0.6510 | 1.0000 | |
| | *S. pistillata* | 42 | control - fibers | 1.7532 | 7.6890 | 0.2280 | 1.0000 | lmer, healthy ~ treat + (1 \| col) + (1 \| origin) |
| | | | control - tirewear | 7.4290 | 7.6890 | 0.9660 | 1.0000 | |
| | | | control - beach | 7.6073 | 7.6890 | 0.9890 | 1.0000 | |
| | | | control - PE | -0.4597 | 7.6890 | -0.0600 | 1.0000 | |

36
